# Reactivation of reward representations associated with the reinforcement of behavior

**DOI:** 10.1101/2024.12.23.628672

**Authors:** Ai Phuong S. Tong, Vishnu Sreekumar, Sara K. Inati, Kareem A. Zaghloul, Mark W. Woolrich

## Abstract

Reward-related representations are found distributed throughout many human subcortical and neocortical regions that support different neural processes. These representations get reinstated for different but related tasks. There is rising evidence that representations reactivate during learning to reinforce prior rewarding choices. However, there remains a critical lack of understanding for whether and how reactivations in humans can facilitate behavioral reinforcement. To investigate this, we recorded from the temporal lobe and prefrontal cortex with intracranial electrocorticography while human subjects learnt two-choice decisions across two different scenes. Crucially, subjects were able to straightforwardly reactivate knowledge gained in a prior scene to make decisions in the current scene, but only when the reward contingencies stayed the same between the two scenes. Using a Bayesian learner, we inferred reward expectation from choice behavior, and measured representations of these reward expectations in electrocorticography data. We found that the representations of reward expectation in the medial temporal lobe and orbitofrontal cortex were reactivated, only when the subjects could straightforwardly transfer knowledge between the two scenes. In the anterior temporal lobe, the reactivation of reward representations increased as learning increased, suggesting an area of plasticity during learning. This work presents a novel framework for measuring the reinforcement, or reactivation, of representations from human electrophysiology during the reinforcement of behavior.

**Significance statement:** This study provides a framework for how the brain processes and reinstates reward-related information during the reinforcement of behavior. Using intracranial electrocorticography, the researchers found that reward information is encoded in distant brain regions, the medial temporal lobe and orbitofrontal cortex. These representations are dynamically reinstated across scenes when reward contingencies align and when knowledge can be transferred. Additionally, reward-related representations in the anterior temporal lobe become increasingly reinforced or reinstated as learning progresses, revealing an area of plasticity during learning. These findings offer new insights into the neural mechanisms underlying decision-making and highlight potential avenues for addressing conditions like addiction or depression, where reward processing and decision-making are disrupted.

## Introduction

Complex behaviors rely on structured representations of information.^1–3^ Representations in the prefrontal cortex have been shown to demonstrate this structure through the mixed selectivity of neurons that display adaptive coding and exhibit highly diverse responses that change over time.^4,5^ Importantly, the same representations can be reinstated in related situations. For example, the overlap in responses of neurons to variables have been shown to be common between experiences that are linked by time, item, or context.^6–9^ In the orbitofrontal cortex, signals encoding task-relevant information are maintained across problems when prior experience facilitates learning.^9–11^ Humans and animals also benefit from representations of abstract information, such as reward expectations.^12^ Representations of abstract task information have been found to be reinstated across different items or contexts.^13,14^

There is evidence that reward-related representations can be found in many distributed regions of subcortex and neocortex, including the orbitofrontal, ventromedial and dorsolateral prefrontal, cingulate, and parietal cortex.^15–24^ While there is good evidence that reward-related representations are found in different parts of the cortex, there is limited knowledge of when and how these representations get learnt and reactivated.

Here, we investigated the capacity of the medial temporal lobe and neocortex to reactivate representations of reward expectation. To do this, we recorded simultaneous electrocorticography (ECoG) in temporal and prefrontal cortices while human subjects performed a task in which they learnt two-choice decisions across different scenes. The subjects were patients with refractory epilepsy undergoing invasive electrophysiological monitoring for seizure localization. The scenes corresponded to different spatial environments, or contexts, within which the choices were presented. Importantly, the task was designed such that subjects were sometimes able to straightforwardly transfer knowledge between scenes because the reward contingencies stayed the same between scene 1 and scene 2, while other times they could not.

By using a Bayesian learner to statistically infer expectation for reward as it varies over trials in human choice behavior, we were able to measure representations of reward expectation in the ECoG data. We measured the similarity of representations of reward expectation between the different scenes and computed the contribution of each brain region to the similarity. This allowed us to investigate where in the brain representations are reactivated and in which scenarios. In particular, we expected representations to be reactivated more when the task allowed information to be transferred between scenes versus when it could not. Finally, we explored the way in which the representations changed with learning.

## Results

The advantage of a reusable representation is it facilitates learning when structure is maintained and relevant across different problems.^14,25–35^ A lack of a reusable representation, resembling a model-free strategy, can be limiting and inflexible because of the need to learn from scratch. This has previously been seen, for example, when there are changes in the environment.^36^

### Task and behavior

We trained human subjects using a behavioral paradigm with a learning (or encoding) phase and a decision (or retrieval) phase with simultaneous electrocorticography (ECoG) recordings (Fig. 1a). We focused on the learning phase in which subjects learned the most rewarding choice between two alternative items based on reward feedback (Fig. 1a). Subjects were trained through a series of “blocks” with instructions about the task shown at the beginning of each scene (see later) in each block. Human subjects were tasked with learning the most rewarding choices of two different “item sets”, where each item set consists of two items that the subject must choose between, and whose reward probabilities are coupled such that the probability of a reward over the two items sums to one. One item set corresponded to a choice between a building versus a face, and the other item set corresponded to a choice between an item of cutlery versus an animal (Fig. 1b). For example, in the building versus face item set, if the faces (for example) are associated with a reward 80% of the time, then the buildings are associated with a reward 20% of the time.

**Figure 1.**
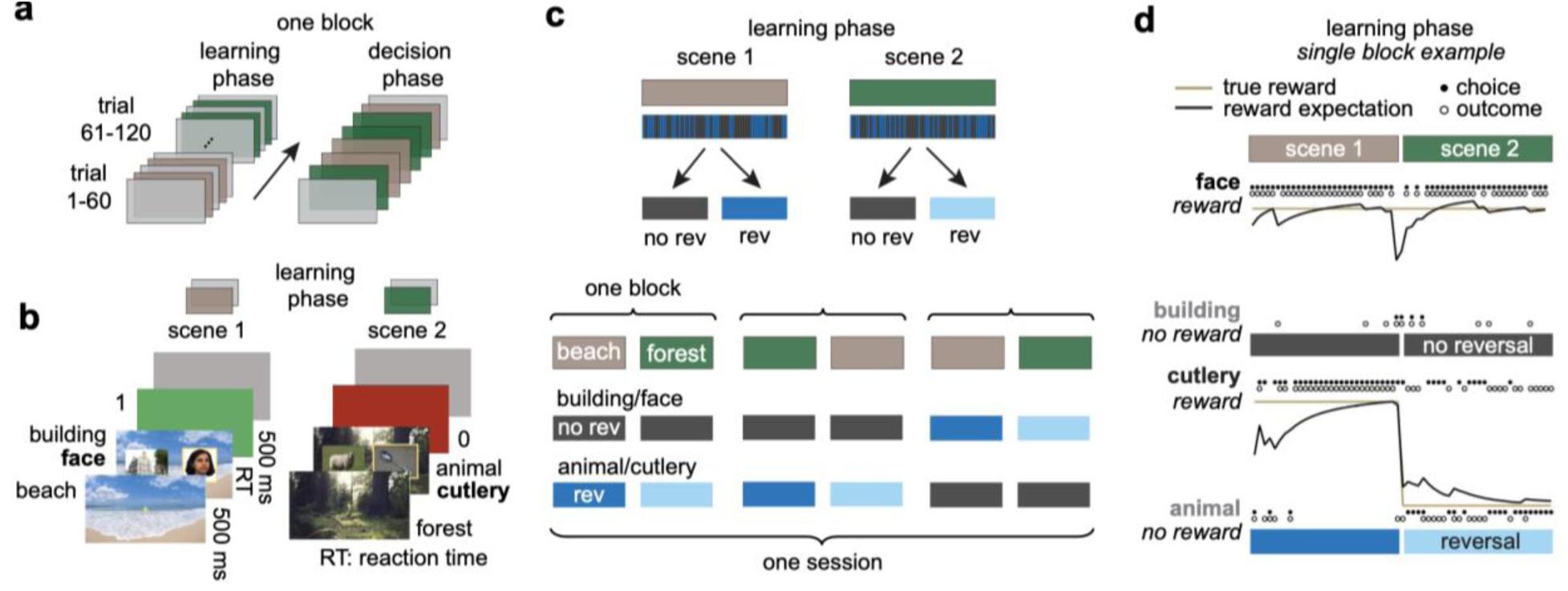
Probabilistic reward learning task modulates reward expectation. **a**, Learning and decision phases of a two-choice decision making task with reversal learning between scene 1 (tan) and scene 2 (green) for an example block. Within a block is a learning phase that consists of 120 trials where reward feedback is given for choices. Following the learning phase is the decision phase where subjects make choices and are not given reward feedback. In this work we focused on the learning phase in which reward feedback is given for choices to investigate representations of reward expectation during learning. **b**, Single trial examples of alternative options in a beach context and then forest context. Subjects are first placed into a scene context for 500 ms and then the item set is presented as two items side-by-side within the scene for subjects to choose between. Based on the subject’s choice, they are given feedback, a reward (green screen) or no reward (red). The images of faces used in this study came from the Face Place dataset.^76^ **c**, [Top] separation of reward reversal (rev) and no-reversal (no rev) trials from scene 1 and scene 2 into four different subsets of trial types, or “context conditions.” The four context conditions correspond to: (1) scene 1 trials for the reversal item set; (2) scene 1 trials for the no-reversal item set; (3) same as condition 1 but for scene 2; (3) same as condition 2 but for scene 2. [Bottom] examples of item sets within the four context conditions in each block over three exemplar blocks in a session. Note, there are two item sets used: 1) building vs face and 2) animal vs cutlery. **d**, Trial course for choices with no reward reversal (top) and with reversal (bottom) between scene 1 and scene 2 for one example block. Choice is indicated by a filled circle and reward outcome by an open circle. Reward expectation, obtained using a Bayesian learner trained on the reward history (see fig. S1), at each trial inferred for face and animal choice shown in black. True reward rate shown in gold.

Subjects were tasked with learning the high reward probability item in two different scene contexts, a beach and a forest. In each block, either beach or forest was selected as the scene context for the first 60 trials, which we refer to as “scene 1”; then the second 60 trials were carried in the other scene context, which we refer to as “scene 2.” Subjects were instructed that there was a reward reversal between the two different scenes in a block for one item set, but no reward reversal for the other.

As an example, consider the case where the cutlery versus animal item set is in the reversal condition. This would mean that the face versus building item set must be in the no-reversal condition. In addition, if we consider the case where cutlery is most frequently rewarded in the beach context, then since the reward contingencies switch for this item set, animals will become the most frequently rewarded in the forest context. Finally, if we consider the case where faces are most frequently reward in the beach context, then since the reward contingencies do not switch for this item set, faces will remain the most frequently rewarded in the forest context as well. Note that for the item set in the reversal condition, the scene context is relevant to the task since the most rewarding choice depends on the scene; whereas for the item set in the no-reversal condition, the scene context is not relevant to the task.

In what follows, we will refer to four different subsets of trial types as “context conditions,” which correspond to: (1) trials from scene 1 (i.e. the first 60 trials of a block) for the item set with a reward reversal, (2) trials from scene 1 for the item set without a reversal, (3) trials from scene 2 (i.e. the last 60 trials of a block) for the item set with a reward reversal, and (4) trials from scene 2 for the item set without a reversal. Also, we will refer to “trial timepoints” as the time points from two seconds before, to one second after, the choice is made for each trial.

Each block consists of a sequence of trials for which the scene context changed from the first 60 trials to the second 60 trials, but the meaning of the scene contexts was fixed for each item set. In other words, the item set in the no-reversal condition and the item set in the reversal condition is fixed within a block. The most frequently rewarded item in the no reversal item set is fixed in 30 trials in scene 1 and in 30 trials in scene 2 within a block, and the most frequently rewarded item in the other item set is one item in the set for 30 trials in scene 1 and the alternative item in the set for 30 trials in scene 2 within a block (Fig. 1c). One of the two item sets are randomly selected to be presented at each trial. Over different blocks, the meaning of the scene contexts was changed, as was the most frequently rewarded item in each item set (this was randomized).

Overall, we trained 17 human subjects and excluded 6 who did not meet at least 70% performance or did not complete at least three experimental blocks in a recording session, which is composed of three to six consecutive blocks that are all completed within one day. The 11 remaining subjects were trained over 3.9 (+/− .1, [3-5]) blocks on average (+/− s.e.m. [range]) over 2.4 (+/− .5, [1-6]) sessions. Each session is performed on a different day.

We aimed to use the behavioral paradigm described above to assess the extent to which representations of task-related information are “reactivated” in related situations. Specifically, in the “no reversal” item set, the most frequently rewarded item is *fixed* between scene 1 and in scene 2, and so we would expect that subjects could transfer knowledge (i.e. reactivate representations) between the two scenes. Whereas, in the “reversal” item set, the most frequently rewarded item *changes* between scene 1 and in scene 2, and so we would expect that subjects would find it harder to transfer knowledge between the two scenes.

### Modulation of reward expectation and reaction times during learning

In an environment where reward outcomes are stochastic or change, an agent dynamically learns to adapt the rate of learning with choices.^37–41^ Thus, we modeled the prediction of possible reward with a Bayesian framework, where the probability of reward for each trial is learnt as a posterior distribution centered around an expected reward probability (fig. S1a-b). With our Bayesian learner, we assumed that this distribution is updated using feedback obtained as a result of each choice that is made. The Bayesian learner learns the distribution of reward probability separately for each item set and for each block, and does not include knowledge of the reversal and no-reversal of reward for item sets. This is because, empirically, we did not find any significant differences in the average probability of choosing the high reward item between the context conditions.

The Bayesian learner is used to update what the subject expects to be the reward probability for an item (i.e. the mean of the posterior distribution of the reward probability) at each trial, using the sequence of choices and reward outcomes. The expected reward probability ranges from 0 to 1, where a value of 1 indicates that a reward for the corresponding item is expected 100% of the time (Fig. 1d). For example, when a reward is not given for choosing the most frequently rewarded item on a later trial, the expectation drops minimally compared to before in an earlier trial (Fig. 1d).

We tracked learning at each trial using the absolute difference between the expectation for reward and the true reward probability, i.e. the absolute value of the reward expectation error (Fig. 2a). Subjects that demonstrated learning formed an expectation for reward that had a low reward expectation error. Within a block, subjects demonstrated a change in the total reward expectation error (i.e. the sum of the reward expectation error across trials) from scene 1 to scene 2 (Fig. 2b). The change in the total reward expectation error varied based on whether there was a reversal in reward between scene 1 and scene 2 and varied over subsequent tasks over blocks. Subjects that demonstrated more efficient learning had a greater decrease in reward expectation error over blocks.

**Figure 2.**
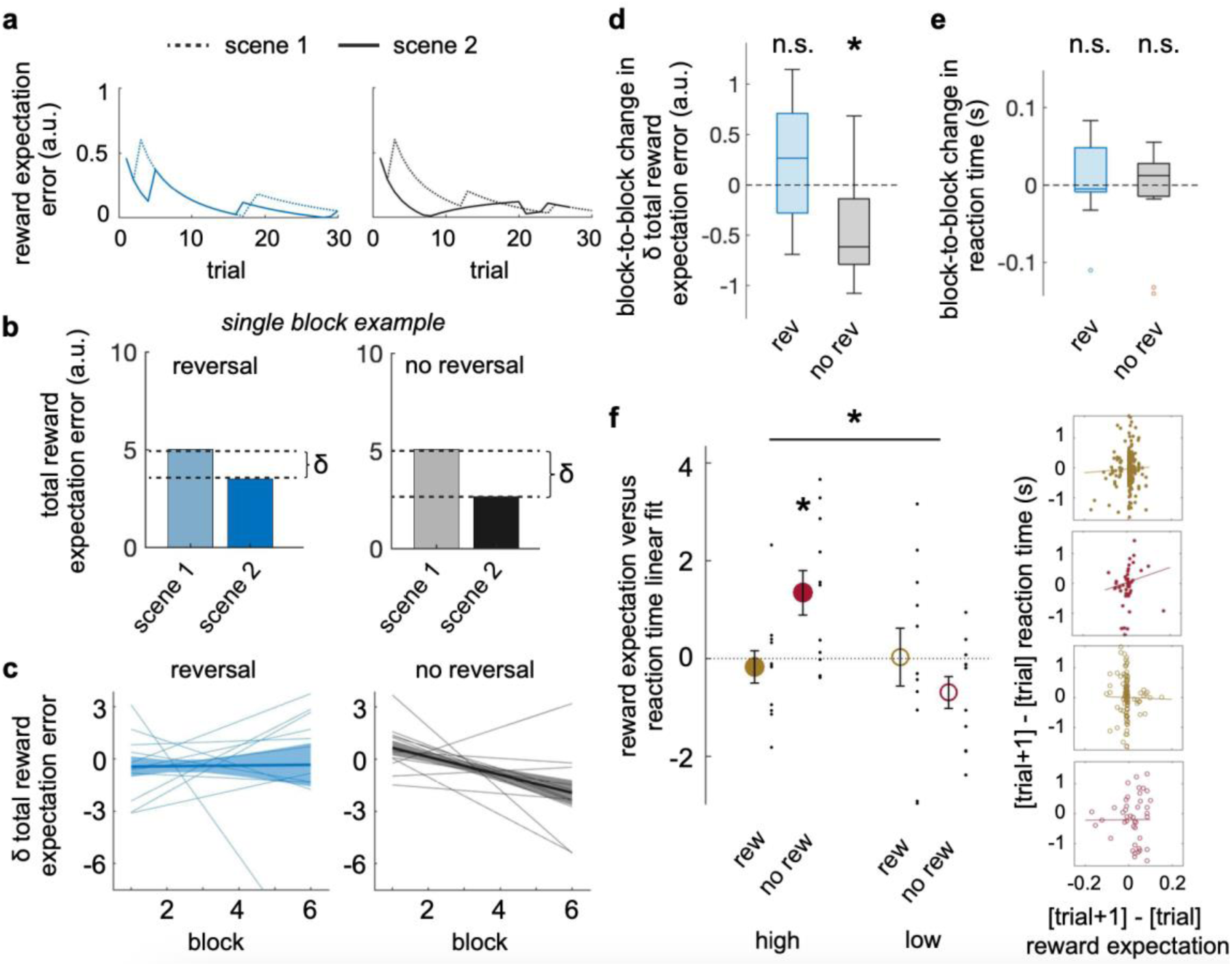
Reward expectations improve with learning and are associated with reaction times. **a**, Single block example of reward expectation error, i.e. the difference between the reward expectation and true reward rate, for reward reversal and no reversal in scene 1 and scene 2. Note that the reward expectations are inferred using a Bayesian learner trained on the reward history (see fig. S1). **b**, Single block example of total sum of reward expectation error for each of the four context conditions. **c**, Measure of block-to-block change in total (summed over all relevant trials) reward expectation error between scene 1 and scene 2, which we refer to as “δ total reward expectation error,” estimated from a mixed effects linear regression for reward reversal and no reversal. Each line reflects a subject, modeled over a minimum of three blocks, on average 3.9 +/− .1 [3-5] blocks over 2.4 +/− .5 [1-6] sessions. Note that δ total reward expectation error can be thought of as the extent to which participants have learnt the presence of the no-reversal and reversal reward structure in the task. **d**, Box plots of rate of change in the δ total reward expectation over blocks for reward reversal and no reversal (mixed effects across-subject t-test, rev, t(10) = .059, p = .954; no rev, t(10) = –2.589, p = .027). **e**, Box plots of rate of change in the δ mean reaction time between scene 1 and scene 2 over blocks for reward reversal and no reversal (across subject two-tailed t-test against zero, *p < .05). As with the total reward expectation error, “δ” refers to the change between scene 1 and scene in two. **f**, Linear fit between change in reward expectation and reaction time for trials after reward (rew) or no reward (no rew) for high and low reward probability choices (left, one-way ANOVA, F(1,40) = 6.54, p = .014; across subject two-tailed t-test against zero, high/no rew, t(10) = 2.98, p = .014). Trial-by-trial relation, where each dot is a trial, pooled across subjects, with the line showing the line of best fit between the difference in reward expectation and difference in reaction time (right plots).

We hypothesized that subjects would be able to learn more efficiently in the context conditions in which knowledge can be straightforwardly transferred between scene 1 and scene 2, i.e. the no reversal item set. To examine the change in learning efficiency, we correlated the change in total reward expectation error with the block number using a mixed effects linear regression model. With each task over blocks, the reward expectation error progressively improved or decreased, indicating more efficient learning, when there was no reward reversal and not when there was a reward reversal (Fig. 2c,d; mixed effects across-subject t-test, rev, t(10) = .059, p = .954; no rev, t(10) = –2.589, p = .027).

In the non-reversal condition choices can become “habitual” since the same items have been reinforced over both scenes. The reduction in reward expectation, therefore, could be from making habitual choices, which would be contrary to the idea that subjects were computing reward expectations. To assess whether choices were habitual, we assessed whether reaction times (RT) similarly decrease over blocks. If choices were becoming habitual, we would expect a decreasing reaction time over blocks. There was no significant decrease in reaction time over blocks (Fig. 2e).

The reaction time provides another measure of changes in choice behavior with learning. To further assess the effect of reward outcomes on choice behavior, we further examined whether reward expectations influenced choice behavior by measuring how RTs prolonged after unexpected omission of reward. We performed a mixed effects linear regression between the trial-wise change in reward expectation and change in reaction time after reward (“rew”) or no reward (“no rew”) for choices with a high and low reward probability (“high” and “low”). Subjects had significantly longer RTs when there was a greater change in expectation after no reward for a frequently rewarded choice (high/no rew) than after a reward for a frequently rewarded choice (high/rew) (Fig. 2f; one-way ANOVA, F(1,40) = 6.54 p = .014; across subject t-test, high/no rew, t(10) = 2.98, p = .014). The significant modulation of RTs related to expectation again suggests that reward expectations were related to choice behavior.

Our findings indicated that subjects adaptively learned from choices and the reward received, so we next investigated how this reward information was represented in the brain during learning.

### Representation of reward expectation in the MTL and neocortex

We have shown behaviorally that in the task subjects used reward expectations and with learning formed reward expectations that better matched the true reward differences. Based on prior studies that assessed behavior guided by rules, we hypothesized that when reward information is relevant to the task it should be actively represented in a way that can influence choices and learning.^42–46^ We next examined whether reward expectation, as it varies over trials, is represented in the brain and in which brain regions.

In what follows, we take reward expectation to correspond to the expected reward probabilities for an item set obtained from a Bayesian learner applied to each subject’s choice and reward outcome history. Note that since the probabilities of the two categories in an item set (e.g. animals versus cutlery) are tied together to sum to one, then changes in expected reward probability are proportional to changes in the difference in the expected reward probability for the two options.

We measured how much trial-to-trial variation in neural power could be explained by trial-to-trial variations in reward expectation using a trial-wise linear regression (Fig. 3a-c). We focused on neural power in the high-frequency range (30-80 Hz and 80-120 Hz) given these ranges have been shown to reflect local spiking activity across the cortex.^47^

**Figure 3.**
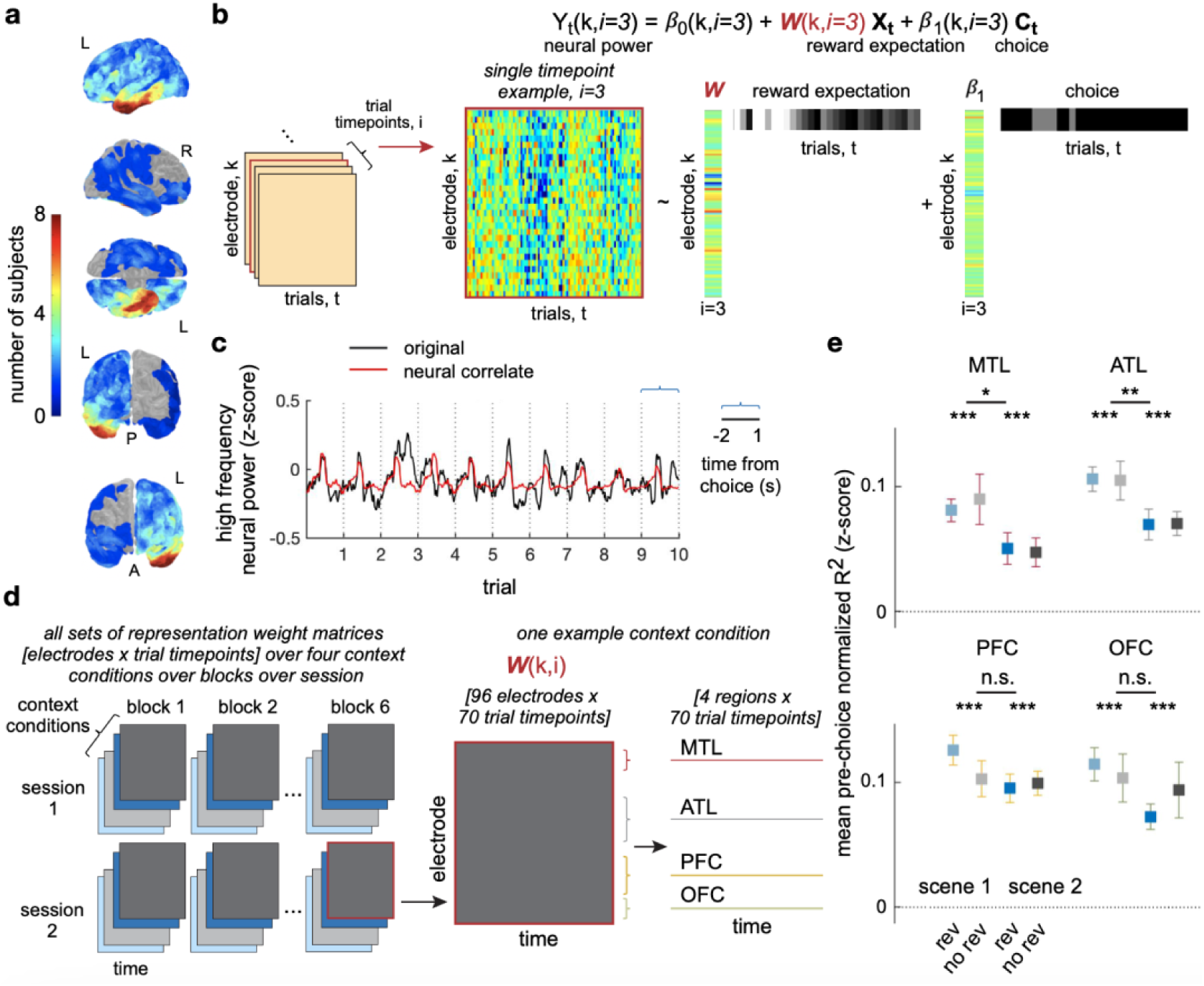
Representations of reward expectation are found in multiple areas distributed across the brain. **a**, Brain plots of electrode coverage across all subjects. L, left. R, right. P, posterior, A, anterior. **b**, Example design matrix of linear regression of neural power, Y(k,i), over trials, t, with expectation, X, and choices, C, as covariates. Trials are organized into subsets of 30 trials organized by context condition. The linear regression is performed over 30 trials for each context condition separately for a single trial timepoint, i, for a single electrode, k, at a time. **c**, Concatenated time series of high frequency neural power (original, black) of the initial 10 trials and concatenated time series of neural correlates of reward expectation (red) of the initial 10 trials from 2.0 sec before to 1.0 sec after the choice is made for a single electrode. **d**, The result of the trial-wise regressions is an [electrodes x trial-timepoints] representation weight matrix, W, capturing how reward expectation is spatially represented at each trial-timepoint [middle]. A version of this matrix is estimated for each context condition (stacked rectangles), block (columns of stacks) and sessions (rows of stacks) [left]. Note that context condition refers to four different subsets of trial types, with each subset corresponding to a combination of reward reversal versus no reversal, and scene 1 versus scene 2. To obtain the extent to which reward expectation is represented in a brain region at each trial-timepoint, we averaged over electrodes within that region, over the trial timepoints within 1.0 sec prior to when the choice is made and over all blocks/sessions/subjects [right]. **e**, The coefficient of determination, or the variance in neural power that is explained in the representations, across brain regions for each context condition. Each dot is a session. Error bars show the s.e.m. across sessions (significance of normalized R^2^ for each region measured by a two-tailed t-test against zero, ***p < .001; one-way ANOVA comparison between scenes; MTL, F(47,1) = 6.97, p = .011; ATL, F(83,1) = 8.76, p = .0040; PFC, F(67,1) = .55, p = .46; OFC, F(75,1) = 3.79, p = .055). Note that R^2^ can be thought of as the extent to which the regression model captured the variations in neural power, which were found to be similar, ∼10%, across all brain regions across all context conditions (***p < .001).

For each subject, we fit a separate trial-wise regression to each trial timepoint of each context condition of each block of each session for each electrode. Performing each of these regressions *separately* avoids the need to make any assumptions about how the representation of reward expectation varies over trial timepoints, or about how it varies across context conditions and brain regions. Note that behaviorally we found the Pearson’s correlation between reward expectation and the chosen item to be greater than zero. Hence, in the trial-wise regression, we included the chosen item category as a co-regressor, so that variations due to choice could be accounted for (fig. S1). This allows us to isolate how much variation in neural power could uniquely be explained by reward expectation above and beyond choice (Fig. 3b).

The result of one of the trial-wise regressions is a reward expectation regression parameter, W(k,i), which tells us the extent to which variability over trials in high frequency power can be explained by reward expectation at trial-timepoint i and electrode k. If we combine these regression parameters over electrodes and trial-timepoints then we get a [#electrodes x #trial-timepoints] matrix, W, capturing the spatial representation of reward expectation at each trial-timepoint (Fig. 3d [middle]). We can obtain versions of this matrix separately for each context condition, block, and session (Fig. 3d [left]). Finally, to estimate the extent to which reward expectation is represented in a brain region, we averaged the representation of reward expectation over electrodes with that region, over the trial timepoints within 1.0 sec prior to when the choice is made, and over all blocks for all sessions and subjects (Fig. 3d [right]).

Using the estimates of the representation of reward expectation within each region, we wanted to know how much the variations in the representations were from variations in neural power related to reward expectation, instead of variations that were predicted by chance. We thus evaluated the regression parameter weights by measuring the “coefficient of determination” (R^2^). This value assesses the extent to which the representations of reward expectations capture the variation in neural power over trials, for each brain region and each context condition (see Methods). The coefficients of determination showed that the same amount of variation in neural power was measured for each context condition, approximately 10%, meaning that the regression model did not fit differently for different context conditions. This suggested that differences in regression parameter weights were not due to differences in how well the model fit to the data (Fig. 3e). When we performed a cross-validation analysis to determine if the regression model for one context condition could explain variations in another context condition, we found low explanatory power (fig. S4). These results collectively demonstrate that across multiple regions of the brain, variations in neural power reflected reward expectations specific to the context condition and were not observed by chance.

Crucially, for both item sets and both scenes, we found, behaviorally, that the reward expectation gradually plateaued to the true reward rate. To validate that the regression model was measuring how much variations in neural power explained trial-by-trial variations in neural power related to reward expectation and not general drifts in neural power that was not task-relevant, we measured how much the variations in the representations for an item set in a scene explained variations in neural power related to reward expectation for an alternative item set and alternative scene. The explanatory power was only high for the neural power of the context condition in which the model was trained, and low for the neural power of the other context conditions (fig. S4). These findings suggest that representations of reward expectation were unique to each context condition. If it was not, then the model that was learnt for one context condition would be able to explain variations in neural power in another context condition.

### Similar and separable representations

We have shown that reward expectation is represented across different brain regions distributed across the cortex. However, this did not mean that the representations of reward expectation in each of those brain regions would always be the same, i.e. always be reactivated, at other points in time.

We hypothesized that only those brain regions that represent reward expectation, and integrate that information with the representation of items, would have similar representations of reward expectation. Further, this would only occur when the information about rewards can be straightforwardly transferred between scene 1 and scene 2, i.e. for the no-reversal item set. We hypothesized that this would not occur in the reversal item set, even though the stimuli they see in both scenes are the same, because the reward associated with the stimuli will have changed.

Here, the representation of reward expectation in a particular brain region corresponds to the “electrode maps” for that brain region, which contains the representation weights for each electrode, as previously described in Figure 3.^48,49^ We next computed a measure of similarity of the representation of reward expectation using an approach that combines cosine similarity and electrode/region ablations (see Fig. 4a,b and Methods). Note that this measure is computed for each brain region, context condition and trial-timepoint.

**Figure 4.**
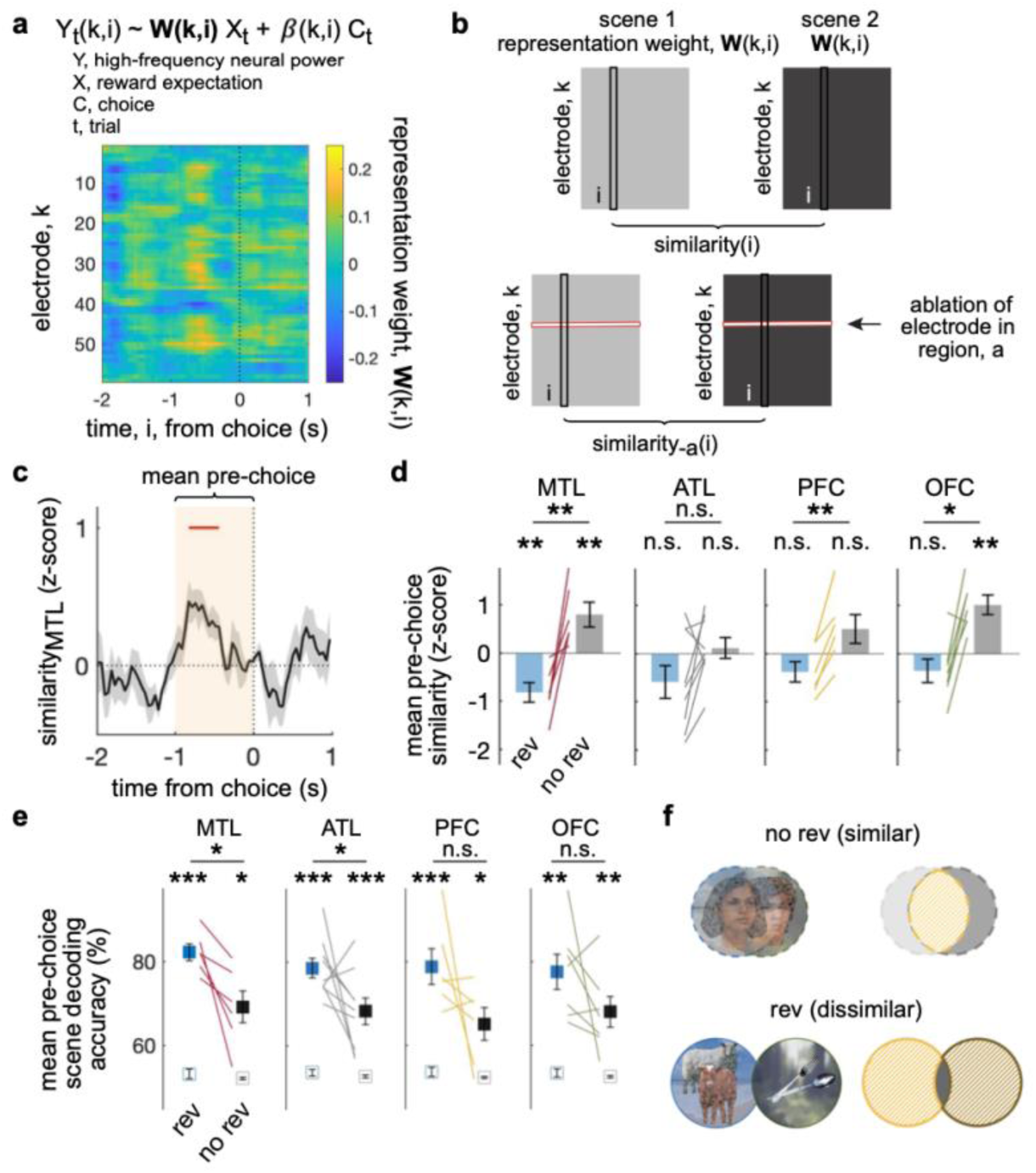
Representations of reward expectation are reusable. **a**, The representations of reward expectation correspond to the reward expectation parameter weights, W(k,i), from the regression of reward expectation onto neural power as they vary over trials, as illustrated in Fig. 3b. **b**, Schematic of how to compute a measure of the contribution of a brain region to the similarity of reward representation between scene 1 and scene 2. First, similarity, for a trial-timepoint, i, is computed using the cosine similarity of “electrode map” that contain the representation weights of reward expectation, W(k,i), for each electrode. Second, the contribution to the similarity for each electrode is computed by ablating each electrode in turn. Finally, the contribution to similarity of each brain region is measured by averaging over the electrodes in that region. **c**, Time course of choice-locked similarity of representations of reward expectation between scene 1 and 2. Shown for the medial temporal lobe (MTL) as an example. Mean and s.e.m. across subjects shown by black shading. Red bar, p<.01, cluster corrected. **d**, [Left] Mean pre-choice similarity measure for MTL, obtained by averaging the pre-choice similarity in©) over the shaded time window. Each line is a subject. [Right] Mean pre-choice similarity shown for anterior temporal lobe (ATL), lateral prefrontal cortex (PFC), and orbitofrontal cortex (OFC). Across subject t-test –-MTL: rev, t(4) = –1.42, p = .23; no rev, t(4) = 4.55, p = .010; OFC: rev, t(5) = –2.11, p = .089; no rev, t(5) = 5.79, p = .0022; **p < .01, n.s., not significant, p > .05. **e**, Mean pre-choice scene decoding accuracy (i.e. decoding scene 1 versus scene 2) using the representation weights of reward expectation, W(k,i), as features; obtained by averaging the decoding accuracy over the shaded time window shown (**c**). Filled squares with error bars show the mean +/− s.e.m. across subjects. Open squares show the chance decoding accuracy from shuffled labels. Significance was measured from the across-subject t-test (*p<.05, **p<.01, ***p<.001). **f**, Overall, in contrast to the reversal item sets, the no-reversal item sets show significantly similar or overlapping representations of reward expectation between scene 1 and 2. This suggests that representations of reward expectation are being reactivated between scene 1 and scene 2, but only when knowledge is expected to be straightforwardly shared. The images of faces used in this study came from the Face Place dataset.^76^

We found that in the 1.0 sec prior to when the choice is made, the representation similarity between scene 1 and scene 2 in the medial temporal lobe increased significantly more for the item set for which there was no reversal, compared to the item set for which there was a reversal (Fig. 4c; mean and s.e.m. across subjects in black shading; red bar, p<.01, cluster corrected).

In the medial temporal lobe (MTL) and orbitofrontal cortex (OFC) the representation of reward expectation 1.0 sec prior to when the choice was made showed significantly greater than zero similarity between scene 1 and scene 2 for the item set for which there was no reward reversal, but not for the item set for which there was a reward reversal (Fig. 4d; across subject t-test; MTL: rev, t(5) = –3.94, p = .011; no rev, t(5) = 3.09, p = .027; ATL: rev, t(8) = –1.77, p = .12; no rev, t(8) = .52, p = .62; PFC: rev, t(5) = –1.84, p = .12; no rev, t(5) = 1.71, p = .15; OFC: rev, t(6) = –1.48, p = .19; no rev, t(6) = 4.89, p = .0027; **p<.01, *p<.05, n.s. p>.05). The representations of reward expectation in MTL and OFC were more similar (between scene 1 and 2) for the item set for which there was no reversal, compared to when there was a reward reversal (Fig. 4d; across subject t-test, rev v no rev, MTL: t(5) = 4.27, p = .0079; ATL: t(8) = 2.018, p = .078; PFC: t(5) = 5.21, p = .0034; OFC: t(6) = 4.017, p = .0070; **p<.01, n.s. p>.05).

Figure 4d shows that there was a significantly greater than zero similarity in the reward representations (between scene 1 and 2) for the reward reversal item sets in MTL and OFC. However, this does not mean that the relevant representations in scene 1 and 2 are identical since the representations can still have components that are orthogonal to each other.^50^

A useful, complementary measure is to look at how “separable*”* the representations are for scene 1 and scene 2. A brain area in which there is high separability is one in which the reward representations look very different to each other between scene 1 and scene 2. To measure separability, we calculated how well the two scenes can be discriminated in a decoding model, using the “neural correlates” of reward expectation as input features. The neural correlates of reward expectation refers to the neural power predicted by the regression model in Fig. 3b using the estimated reward expectation parameter weights, W(i,k) (see Methods).

We found that scene 1 versus scene 2 can be decoded more accurately from the neural correlates of reward expectation when the scene context is relevant to the task (i.e. for the item set for which there is a reversal) compared to when the scene context is not relevant (i.e. for the item set for which there is no-reversal). This suggests that representations selectively vary more between scene 1 and scene 2 when there is a reward reversal, compared to no-reversal (Fig. 4e; fig. S7-8).

Overall, in contrast to the reversal item sets, the no-reversal item sets show significant similarity and less different representations of reward expectations between scenes 1 and 2. This suggests that the representations of reward expectation are being reactivated between scene 1 and scene 2, but only when knowledge is expected to be straightforwardly shared. Note that this is only possible if the participants have learnt, and are acting on, the presence of the no-reversal and reversal structure in the task.

### Association of representations with learning

To further probe the behavioral relevance of the representations of reward expectation, we investigated whether the similarity of the representations between scene 1 and scene 2 were associated with learning. Here, we defined “learning” as the extent to which participants have learnt, and are acting on, the presence of the no-reversal and reversal reward structure in the task. To measure this, we estimated how much the sum of the reward expectation error over trials change between scene 1 and scene 2, which we refer to as the “δ total reward expectation error.” Specifically, this refers to the difference in the sum of the reward expectation error across 30 trials in scene 2 minus the sum of the reward expectation error across 30 trials in scene 1 for an item set. We expect learning corresponds to a reduced or negative change, δ, in total reward expectation error from scene 1 to scene 2.

Then, we correlated the δ total reward expectation error between scene 1 and scene 2 with the similarity in representations (cf. Fig. 4d) between scene 1 and scene 2 as both measures varied over blocks. This correlation was measured for each session and then we averaged the correlations across sessions for each subject. A significant positive correlation was found in the ATL, such that as learning increased then so did the similarity between scene 1 and 2, but only for the item set for which there was no-reversal in reward, when knowledge can be straightforwardly transferred between scenes; and not for the item set for which there was a reversal in reward, where knowledge cannot be straightforwardly transferred (Fig. 5a; ATL: rev, t(8) = .99, p = .35; no rev, t(8) = –2.43, p = .041; rev v no rev, t(8) = 4.04, p = .0038; n.s. not significant, p>.05).

**Figure 5.**
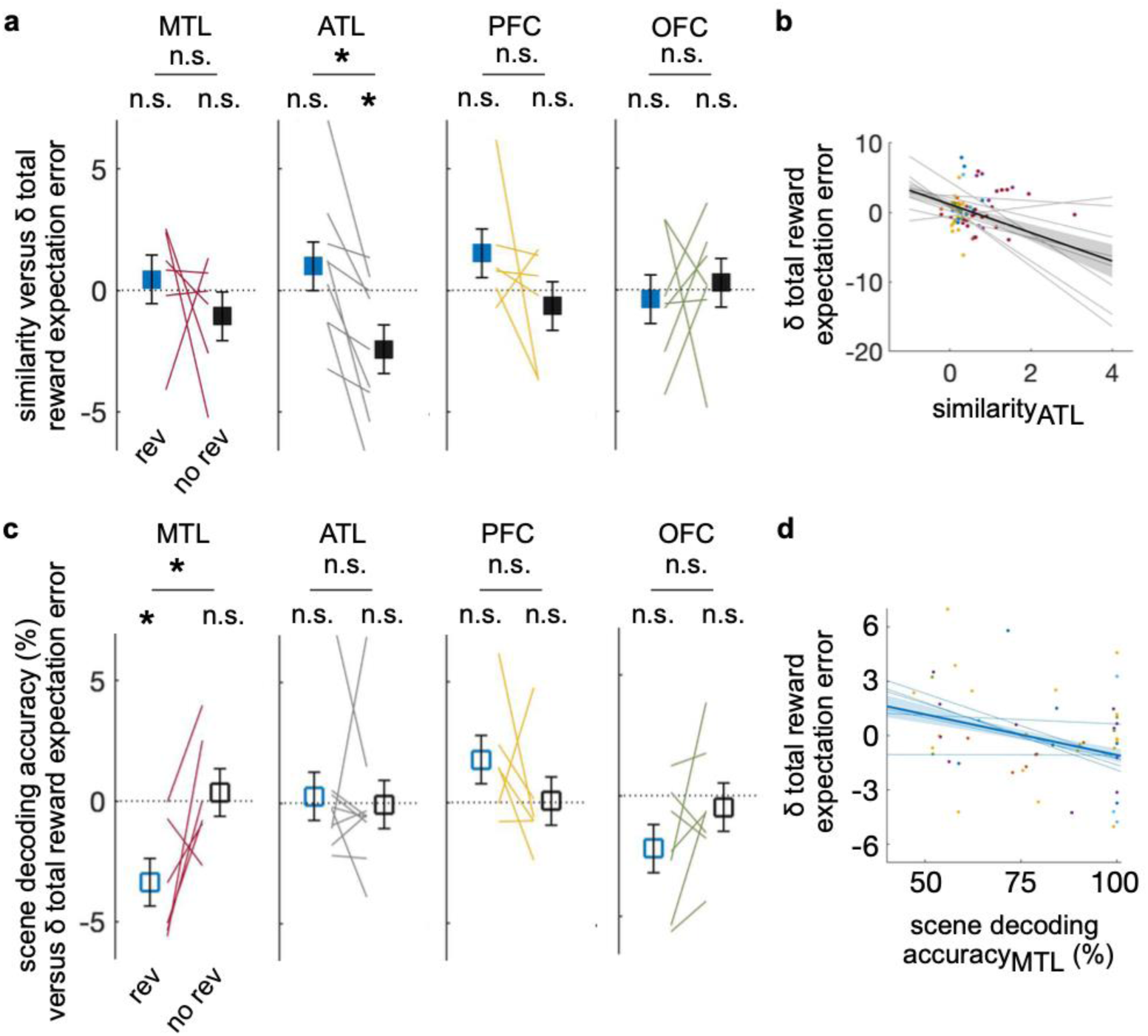
Similarity of representations are associated with learning performance. **a**, Association between the block-wise measures of representation similarity (cf. Fig. 4d) and of change in δ total reward expectation error between scene 1 and scene 2 (mixed effects linear regression; across-subject t-test ATL: rev, t(8) = .99, p = .35; no rev, t(8) = –2.43, p = .041; rev v no rev, t(8) = 4.04, p = .0038; n.s. not significant, p > .05; **p < .01, *p < .05). Note, the representation similarity can be thought of as the extent to which representations of reward expectation are being reactivated and, as before, δ total reward expectation can be thought of as the extent to which participants have learnt the presence of the no-reversal and reversal reward structure in the task. **b**, Relation between the similarity of ATL representations (cf. Fig. 4d) and changes in expectation error between scene 1 and scene 2. Each dot is a block, pooled across sessions, and each color is a subject. Each thin line is the linear fit for a subject. Thick black line and shaded area is the mean +/− s.e.m. of fits across subjects. **c**, Association between scene decoding accuracy using reward representations (cf. Fig. 4e) and change in δ total reward expectation error between scene 1 and scene 2 (mixed effects linear regression; across-subject two-tailed t-test, MTL: rev, t(5) = –3.37, p = .020; no rev, t(5) = .35, p = .74; rev v no rev, t(5) = – 2.72, p = .042, *p < .05). **d**, Relation between scene decoding accuracy of MTL representations and changes in reward expectation error between scene 1 and scene 2. Each dot is a block, pooled across sessions, and each color is a subject. Each thin line is the linear fit for a subject. Thick blue line and shaded area is the mean +/− s.e.m. of fits across subjects (*p < .05).

We next correlated the δ total reward expectation error between scene 1 and scene 2 with the ability to discriminate between scene 1 and scene 2 (cf. Fig. 4e) as both measures varied over blocks. A significant positive correlation was found in the MTL, such that as learning increased the ability to discriminate between the two scenes also increased, but only for the item set for which there was a reversal in reward, when knowledge cannot straightforwardly be transferred between contexts (Fig. 5b; MTL: rev, t(5) = –3.37, p = .020; no rev, t(5) = .35, p = .74; rev v no rev, t(5) = –2.72, p = .042).

## Discussion

These findings suggest that the representations of reward expectation overlap and can be transferred when knowledge can be shared straightforwardly between two different contexts. The overlap in the representations between context conditions changed adaptively over time with learning, indicating that a potential basis for learning to learn efficiently relies on forming sets of reward-related representations that can be reactivated.

Our results contribute to understanding the distribution and reactivation of reward representations during learning in several important ways. Using ECoG in human subjects who performed a reversal learning task, we measured representations of reward expectation and observed these distributed patterns of activity re-appear over related situations between two different scenes (Fig. 4). Importantly, when we measured the representation of reward expectation across many brain regions, we see significant similarity in representations between the two scenes across specific not all brain regions. The two association cortices (ATL and dlPFC) do not show reusable or similar representations. Our study is the first demonstration to our knowledge of the learning and use of distributed reward-related representations in humans during a reversal learning task using ECoG recordings over time.

Our findings add to earlier results of similarity in covariance structure,^51^ amplitude of activity or firing rate,^13,52^ neural subspaces,^53^ and neural trajectories^54^ across related but different tasks. Our findings also suggest distributed reward circuits where the representation of reward expectation is integrated into the representation of items across human neocortex as reward is associated with items.

### Spatial generalization

Activation or co-activation of neural activity that is maintained or reactivated across different environments have been described as generalization.^13,14^ Several studies have demonstrated such generalization across task structure,^33^ concepts,^32^ value,^30,31^ experiences,^28^ relational information,^27,35^ shared goals,^34^ and error and conflict.^55^ Our findings add to the neural evidence for generalization of reward information across space. The representation of expectation was ubiquitous across brain regions and repeated transiently in different settings, resembling a retrieval mediated reactivation related to a choice. These findings add to prior studies that have demonstrated distributed reactivation locked to reward outcome,^29^ associated with sensory processing,^56^ and during reinstatement for cued memory retrieval.^49^

### Similarity of representations during learning

Many prior studies revealed that ensembles activated during learning are reactivated based on a range of factors, including being close in time,^6^ depending on task demands,^57^ based on learned rules,^58^ and due to constraints by pre-existing local cortical connectivity that form a manifold of preferred activity patterns.^59,60^ Other experiments describe representations that change in the setting of forming a schema or as a result of learning from prior experience.^9^

We expand on prior results in a crucial way, demonstrating that reward-related representations not only occur in local brain regions to guide learning,^61^ but is integrated into representations in different subcortical and neocortical areas in humans. We also observed similarity in representations in the medial temporal lobe and orbitofrontal cortex at a faster timescale than in the anterior temporal lobe, suggesting that there are distinct systems for using and learning reward-related representations to guide choices.

## Conclusion

Representations that were reactivated were found in medial temporal lobe, consistent with prior studies that showed that learning statistical regularities over time is hippocampal-dependent.^62^ In this study, we assumed that representations are locked to choices and the timing of this locking are the same over different contexts and throughout learning. We would expect changes to occur during learning, for example, with representations forming more rapidly, given we observed faster reaction times over trials.^63^ The relevant representations were rather transient and found within one second of when the choice is made, resembling high frequency events that have been shown to underlie memory-guided processing or reflect the reactivation or reinstatement of prior related events.^64–66^ Ongoing investigations will assess the involvement of memory retrieval processes in the use and strengthening of representations.

## Materials and methods

### Subjects

Eleven subjects (two female, 20 to 57 years-old) with drug-resistant epilepsy underwent a surgical procedure in which platinum recording contacts were implanted on the cortical surface and within the brain parenchyma. In each case, the clinical team determined the placement of the contacts to localize epileptogenic regions. The clinical regions of investigation for the subjects are summarized in Fig. 3 and Table 2.

Data were collected at the Clinical Center at the National Institutes of Health (NIH; Bethesda, MD). The Institutional Review Board (IRB) approved the research protocol (11 N-0051), and informed consent was obtained from the subjects and their guardians. All analyses were performed using custom built Matlab code (Natick, MA). Data are reported as mean ± standard error of the mean (s.e.m.) unless otherwise specified.

Analyses were performed within an experimental block and then results for each session were computed from an average across blocks and results for each subject were computed from an average across sessions. Mixed effects analyses were completed for data pooled over blocks and sessions.

### Behavioral task and analysis (related to Fig. 1 & 2)

#### Task structure

To investigate representations of reward expectation during learning, we designed a two-choice task in which rewards are probabilistic and there is a reward reversal and no reversal between two scene contexts, a beach and forest. There are two pairs of items, or “item sets”, building versus face and cutlery versus animal, that switch pseudorandomly within each scene (Fig. 1b). A block consists of 30 trials of each of these subset of trial types for the two item sets over the two scenes, giving a total of 120 trials in a block. Subjects complete up to six consecutive blocks in a session to repeatedly get exposed to the structure of the task. The context that is first and second is randomized for each block. The item in each item set that has a high probability of reward is randomized for each block. Reward probabilities for the items in an item set sum to one with one item having a higher probability of reward compared to the alternative. The high reward probability is on average 80% (70-90%) across subjects and subsequently the alternative item is rewarded on average 20% (10-30%) of trials. There is some variability in this ratio because if subjects perform at less than 50% in a subsequent decision phase the high reward probability is increased. Once a probability is determined, it is kept constant for all blocks in the experimental session.

Subjects first make choices and receive a reward feedback for their choices over 60 trials in one scene context and then move to a second context and complete another 60 trials. In the second scene, the reward probability for one item set reverses (“reversal” condition) and the other item set does not (“no-reversal” condition).

We instructed subjects to learn to separate the two conditions based on whether reward reverses or does not reverse in a new scene. We assessed how subjects adapted to changes in reward and scene compared to when there is no change in reward and only a change in scene. Subjects completed between three and five blocks. Within each scene, there are 30 trials of each item set, which are interspersed pseudorandomly between the two item sets (Fig 1). This comprises the learning phase of each block and is the focus of this study. Each learning phase is followed by a decision phase, which is not investigated here. Note, during the decision phase that follows the learning phase subjects are not given feedback for choices and must retrieve from memory the rewarding choices that they learned. During this decision phase, subjects are first cued with the scene and then are presented with a pair of items. Thus, over blocks, subjects may learn to increasingly associate the scene with rewarding items to separate what item set has a reversal or no-reversal in reward.

#### Inferring expectation (Bayesian model) (Fig. 1, fig. S1)

Both items in an item set are presented simultaneously at each trial. The probability of reward for the two options are related, where the probability of reward sum to one.

Therefore, we expected that choices are guided based on the difference in reward expectation between the items with the high and low reward probability. Note we do not take the difference for the item that is chosen versus the item that is not chosen because previous studies have suggested that the frame of reference for comparing what is rewarding when two items are presented simultaneously is in the option space (i.e. high minus low reward probability) rather than action space (i.e. chosen minus not chosen item reward probability).^67,68^

At each trial we inferred the expectation for reward from the sequence of choices and reward outcomes. We represent the choice over trials with two binary vectors, one vector for each item set, which consists of a value of 1 for a trial when the high reward probability item is chosen and a value of 0 when the low reward probability item is chosen. Similarly, we represent the reward outcome over trials with two binary vectors for each item set separately, with 1 for a trial when the outcome is a reward and with 0 when the outcome is no reward. For the reversal in reward in scene 2, we use a flipped r where there is a high probability for the previously low reward probability in scene 1.

We approximated the posterior distribution for the reward probability at each trial from the prior and likelihood. We describe the likelihood for the observed sequence of choices given the reward probability as follows:

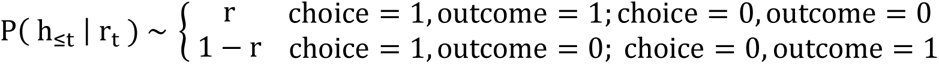

where r is the reward probability distribution and consists of 100 values increasing linearly from 0 to 1, the choice is 1 for the item with the high reward probability and 0 for the item with the low reward probability, the reward outcome is 1 for reward and 0 for no reward, and ℎ_≤*t*_ is the observed history of choices and outcomes up to trial t.

Using this likelihood and a Markovian prior, we computed the posterior distribution over the reward probability as follows:

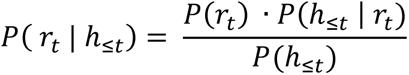

where ℎ_≤*t*_ reflects the history of choices and reward outcomes up to trial t. The prior, P(r_t_), is the previous posterior, P( r_t-1_ | h_≤t-1_). The initial prior is a normalized uniform distribution over the reward probability.

In this way, we measured the reward probability estimate and the uncertainty around it, or the belief of the reward probability, through Bayesian inference with numerical integration. When we normalize the function to integrate to one, the denominator can be disregarded.

We approximated the expectation for reward for an item from the mean of the posterior distribution over the reward probability, which ranged from 0 to 1. We computed the relative reward expectation, or the difference between the mean of the posterior distributions for the high and low reward probability items. Throughout the text, we refer to this relative reward expectation as the “reward expectation.”

#### Reaction time (Fig. 2e,f; fig. S2a)

Reaction time is thought to reflect the cognitive effort needed to change rules because there is a “switch cost.”^2,69^ The question here is do we find a switch cost because reward expectation is accessed when there is no reversal in reward or needs to be recomputed after a reversal in reward? We refer to the sequential sampling decision making literature, in which the accumulation of evidence during decision making is related to reaction times. We can extend the interpretation of reaction times to value-based decision making, where evidence is sampled to predict outcomes or value of items that can be chosen.^70^ Here, we assessed if reaction time is related to reward expectation when reward expectation is used or computed to make a choice. For example, when a subject is not rewarded for an item, such as a face when it has a high probability of reward, if they select the face, then that would mean they computed a higher reward expectation if they select the face in the next trial, despite receiving no reward in the last trial). The hypothesis is that it also would take a longer time for the subject to respond because they need to compute and update the reward expectation.

In order to assess whether subjects compute reward expectation based on feedback for their choices and use the reward expectation to make a choice, we measured the correlation between the change in reward expectation and change in reaction time after an item with a high probability of reward is rewarded or not rewarded and after an item with a low probability of reward is rewarded or not rewarded. This gives us another measure of how much subjects’ choice behavior changes depending on reward expectation. To compare the relation between reward outcomes, we performed an ANOVA that tests for significant differences in the relation between reward expectation after a reward or no reward for a choice and the reaction time.

### Electrophysiological recordings

#### Electrode localization

We collected electrocorticography (ECoG) data from 11 human subjects. Subdural contacts were arranged in both grid and strip configurations with an inter-contact spacing of 10 mm. We captured iEEG signals sampled at 1000 Hz and applied several pre-processing steps to remove movement artifacts and pathological activity related to the patient’s epilepsy, as previously described.^71^

To examine electrodes in the lateral prefrontal cortex, we identified electrodes that map to the frontal pole, rostral middle frontal, and caudal middle frontal parcellations of the Desikan-Killiany atlas. For the medial temporal lobe, we used the entorhinal cortex and parahippocampal gyrus parcellations. For the anterior temporal lobe, we used the temporal pole and superior temporal lobe parcellations. For the orbitofrontal cortex, we identified electrodes that map to the medial and lateral orbitofrontal cortex. For the parietal cortex, we identified electrodes that map to the post central gyrus, supramarginal gyrus, and superior and inferior parietal cortex.

#### Spectral analysis

To measure the neural activity in electrodes, we computed the mean spectral power in bandpass filtered signals from the Hilbert envelope. To account for changes in power across experimental sessions, we z-scored power values separately for each frequency and for each session using the mean and standard deviation for that session. We separately computed the mean 30-80 Hz (high gamma) and 80-120 Hz (ripple band) power in 400 ms trial timepoint windows with 90% overlap from two seconds before to one second after when the choice is made. The narrow 80-120 Hz high frequency band has been associated with ripple events in humans and correlate with synchronous firing of neurons during episodic memory retrieval. ^71–74^

#### Representations of expectation (linear regression) (Fig. 3)

We measured the representation of “reward expectation”, which refers to the relative difference in value for the high versus low reward probability item, by performing a trial-wise regression of the mean power for each electrode and each trial timepoint over the reward expectation. The trial timepoints were 400 ms time windows with 90% overlap from two seconds before to one second after a choice. Note since we performed the regression analysis by individual recording electrode. We do not capture the covariation between recording electrodes through the regression.

Representations can change dynamically over time in each scene for each item set in each block. Thus, we performed a trial-wise regression of neural power on reward expectation separately for each scene and item set in each block to derive a [electrodes x trial-timepoints] representation weight matrix, W(k,i), given by the formula:

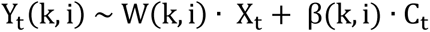

where Y_t_(k,i) is the 30-80 Hz and 80-120 Hz neural power over 30 trials, t, for electrode, k, in a trial timepoint, i. X_t_ is the corresponding reward expectation over the 30 trials, t. C_t_ is the chosen item over 30 trials, t, and is binary: 1 for the item with the high probability of reward and 0 for the item with the low probability of reward. By performing a regression across trials at each trial timepoint, we assumed that representations were similarly locked to when the choice is made across trials.

#### Explained variance of representations (coefficient of determination) (Fig. 3)

To evaluate how well the linear regression model explains the relationship between expectation for reward and neural power, we computed the coefficient of determination (R^2^), which compares the error between the data and the mean of the data and the error between the data and the model prediction.^75^

We computed the sum of squared errors for the regression model, SSE_reg_, and its mean, SSE_tot_ over 30 trials for each recording electrode, k, at each trial timepoint, i.

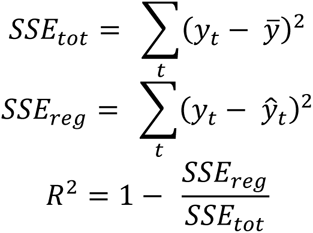

where y is the neural power, 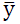 is the neural power mean, and ŷ is the regression-based estimate of neural power. We computed the neural power for 30-80 Hz and for 80-120 Hz separately.

The ratio SSE_reg_ / SSE_tot_ indicates relative error between the model estimate and the variation in data. A ratio greater than one indicates poor model fit to variations in data, and when subtracted from one this results in a low R^2^. A ratio less than one indicates good model fit to variations in data, and when subtracted from one this results in a high R^2^. Thus, the coefficient of determination provides a measure of how much the variance in neural power is captured by the regression model. In order to measure how much the regression model capture variance in neural power that is specific to a context condition, we compared the model estimates to variations in the alternative context conditions and assessed whether task-related trial-by-trial variations are being modeled rather than task-irrelevant drift, which would be present in both scenes and when there is a reversal or no reversal in reward.

For example, we used the regression weights for the representation in scene 1 for the item set with no reversal and computed the estimate of the neural power related to reward expectation for the item set with a reversal in reward (Fig. 3b). To compute the sum of squared errors over the mean, SSE_tot_, for the model trained for the reversal item set in scene 1 and applied it to the no-reversal item set in scene 2, we measured the sum of the squared difference between the neural power for the no-reversal item set in scene 2 and the mean of the neural power in scene 1 for the reversal item set. Then, to compute the sum of squared errors for the regression model, SSE_reg_, we measured the difference between the neural power for the no-reversal item set in scene 2 and the regression-based estimate of the neural power in scene 1 for the reversal item set. We performed this model fit comparison for the alternative reversal condition, alternative scene, and both, for each context condition.

For each context condition in each block, we computed R^2^ for each electrode and each trial timepoint. To compute the R^2^ for each region, we took the mean across electrodes within the region. We computed a mean for each subject by taking the mean over blocks for each session and then the mean over sessions.

#### Neural correlates of expectation (fig. S3)

In order to assess the relation between representations of reward expectation and choice behavior on a trial-by-trial basis, we measured the high frequency neural power predicted by the regression model using the estimated reward expectation parameter weights, which we refer to as the “neural correlate” of reward expectation. We used the reward expectation parameter weights, W(k,i), for each trial timepoint, i, and each electrode, k, and the choice parameter weights, β_1_(*k*, *i*), for each trial timepoint and electrode to estimate the neural power time course of each electrode.

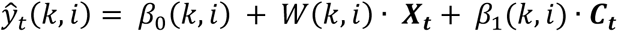

where 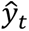 is the 30-80 Hz and 80-120 Hz neural power over 30 trials, t, for electrode, k, in a trial timepoint, i. *X*_*t*_ is the reward expectation over 30 trials, t. *C*_*t*_ is the choice over 30 trials, t.

While we found reward expectation to be represented in multiple areas of cortex, this did not mean that the representation across these areas of cortex would be observed again, or be reactivated, in a similar situation across all cortical areas that represented reward expectation.

We hypothesized that only brain regions that represent reward expectation and integrated this with the representation of items would have similar representations of reward expectation when the knowledge of reward associated with the item can be transferred between scene 1 and scene 2. We found similar representations between scenes localized in two brain regions, not all areas of cortex that we reviewed. We measured representation similarity by the similarity between “electrode maps,” which contain the representation weights for each electrode, during scene 1 and the electrode maps during scene 2.^48^ We assessed for similarity in electrode maps and would expect reactivation of representations when knowledge can be straightforwardly transferred between scene 1 and scene 2.

Crucially, subjects are shown the same stimuli (category of items) in scene 1 compared to scene 2 for the no-reversal and reversal item sets. The difference in the stimuli between scenes comes from the association of these stimuli with reward, which is only different for the reversal item set and not for the no reversal item set. Since we focused on the relation between variations in spectral power and reward expectation prior to when the choice is made, the stimuli that the subjects see between scenes are the same for both the reversal and no reversal item sets.

For the no reversal item set, we would expect the brain responds similarly for the similar stimuli between scenes. For the reversal item set, we hypothesized that the brain response is different between scenes, even though the stimuli they see is the same between scenes, because the brain response changes with the different reward associated with the items.

### Similarity analysis (related to Fig. 4)

#### Electrode and regional representation similarity

We examined the reactivation of representations of individual electrodes by measuring how much each electrode contributes to the similarity of distributed representations between scenes. We used an ablation approach in which we ablate one electrode at a time and re-computed the similarity after each ablation. We take the difference in similarity between the original similarity and the similarity after ablation. A difference greater than zero indicates the electrode is similar between scenes. A difference less than zero indicates the electrode is dissimilar between scenes. To quantify the significance of the similarity between scenes, we compared the empirical similarity to a null distribution of chance similarity. To generate a null distribution, we randomly flipped the scene labels for each electrode and applied a random circular shift in time for each electrode. We again ablated one electrode at a time for each permutation 200 times. To compute the reactivation or similarity of representations in regions, we computed the mean across electrodes in a region.

#### Representation similarity between scenes (Fig. 4d)

To investigate whether representations are observed in both scene contexts, we measured the spatial similarity of representations between scenes. We constructed a representation vector, or electrode map, composed of the reward expectation regression parameter weights, W(k,i), over electrodes, k, at a trial timepoint, i:

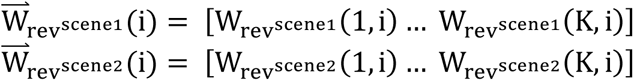

where the [electrodes x 1] regression weight vectors at trial timepoint, i, for the reversal, rev, item set is expressed for each scene separately. This was similarly done for the no-reversal item set.

We computed the cosine similarity of the electrode maps as a function of time, i.

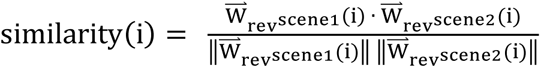

as done previously.^49,65^

To quantify the mean similarity of representations between scenes for each subject, we computed the average across blocks for each session, and then computed the average across sessions. To measure the mean pre-choice similarity, we computed the average in the 1.0 sec prior to when a choice is made.

#### Scene decoding (Fig. 5)

To examine how separable the representations between scene contexts are, we measured the accuracy of decoding scene context from the correlates of expectation. For each block, we trained a support vector machine classifier with 5-fold cross-validation to decode scene using the mean neural correlate of reward expectation in a region at each trial timepoint for an item set in 60 trials, 30 trials in each scene. We measured the mean scene decoding accuracy within one second before the choice is made for each block. We then measured the scene decoding accuracy for each subject by computing the scene decoding accuracy for each session from the mean across blocks and then computed the mean across sessions. We generated the chance decoding accuracy for each subject in a similar way, with random shuffling of the scene labels during the training of the classifier.

## Data availability

Processed recordings and behavioral data will be available online. Raw recordings and behavioral data are available on request.

## Code availability

Code is available at GitHub (https://github.com/tongap3).

## Author contributions

A.P.S.T. contributed to conceptualization, methodology, data curation, formal analysis, writing – original draft, reviewing, and editing. V.S. contributed to data curation, writing – reviewing and editing. S.K.I. contributed to data curation. K.A.Z. contributed to data curation, writing – reviewing and editing. M.W.W. contributed to conceptualization, methodology, supervision, writing – reviewing and editing.

## Materials & Correspondence

Correspondence and material requests should be addressed to Ai Phuong S. Tong and Mark W. Woolrich.

## Acknowledgements

We thank Laurence Hunt, Sanjay Manohar, Ignacio Saez, Ashwin Ramayya, Ben Seymour, Cameron Higgins, Andrew Quinn, Tom Marshall, members of the OHBA Analysis Group lab (Chetan Gohil, Mats van Es), members of the NIH Surgical Neurology Branch (Julio Chapeton, Weizhen “Zane” Xie), and Huiling Tan for feedback and helpful discussions. This project was supported by funding from the National Institutes of Health Oxford-Cambridge Scholars Program and the Goodger and Schorstein Scholarship from Oxford (to A.P.S.T.).

## Supplementary Figures

**figure S1.**
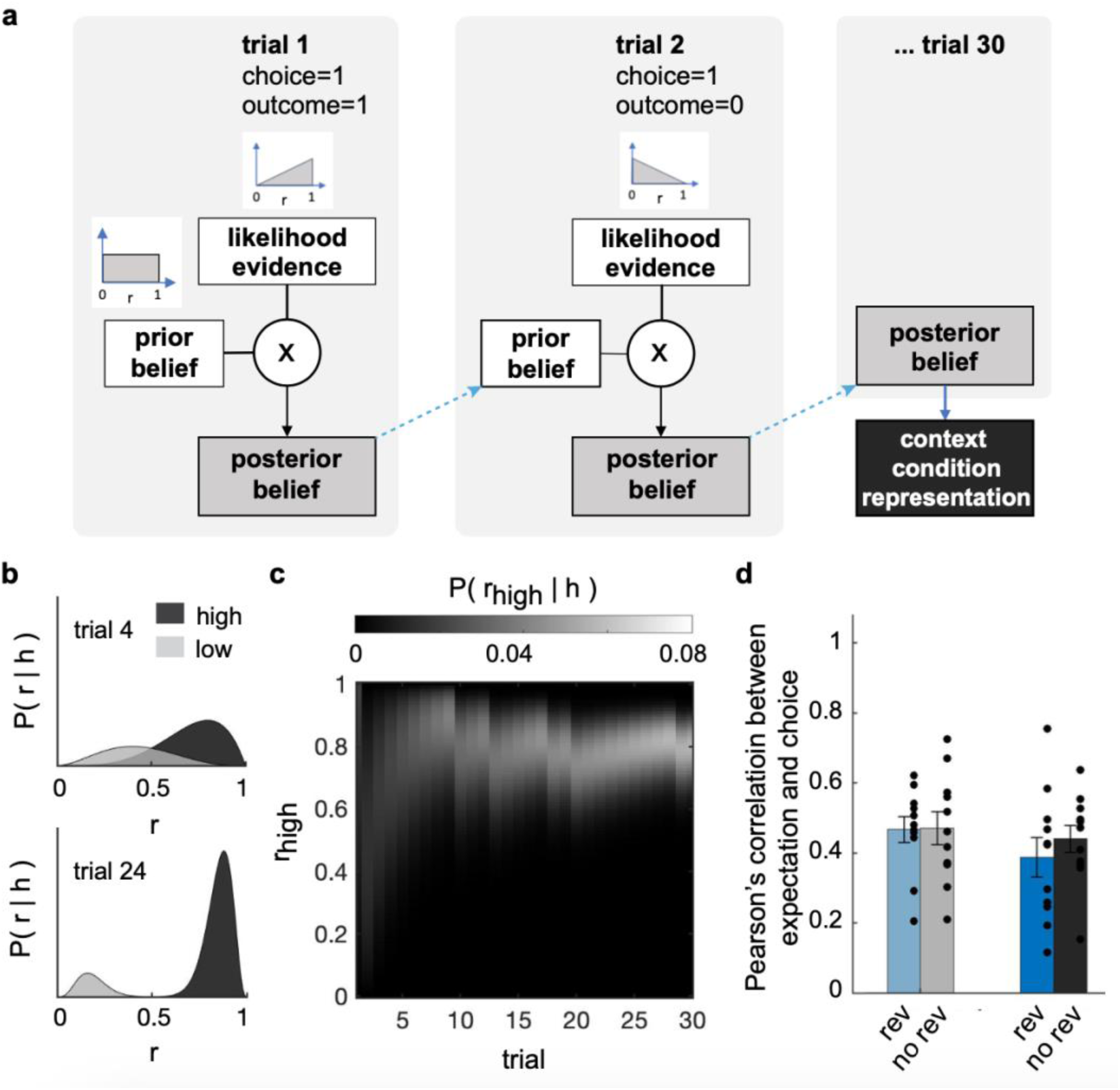
Correlation between expectation and choice. **a**, Schematic of trial-by-trial belief updating process. **b**, Example posterior belief distribution, P(r|h), over the reward probability, r, given the history, h, of choices and reward outcomes (see Methods) for the high reward probability item (black) and the low reward probability item (gray) for an early trial (top, trial 4) and a later trial (bottom, trial 24) within a Scene. **c**, Series of posterior belief distribution, P(r|h), over trials for one example context condition for the high reward probability item. **d**, Pearson’s correlation between the reward expectation and the fraction of choices that are the high reward probability item.

**figure S2.**
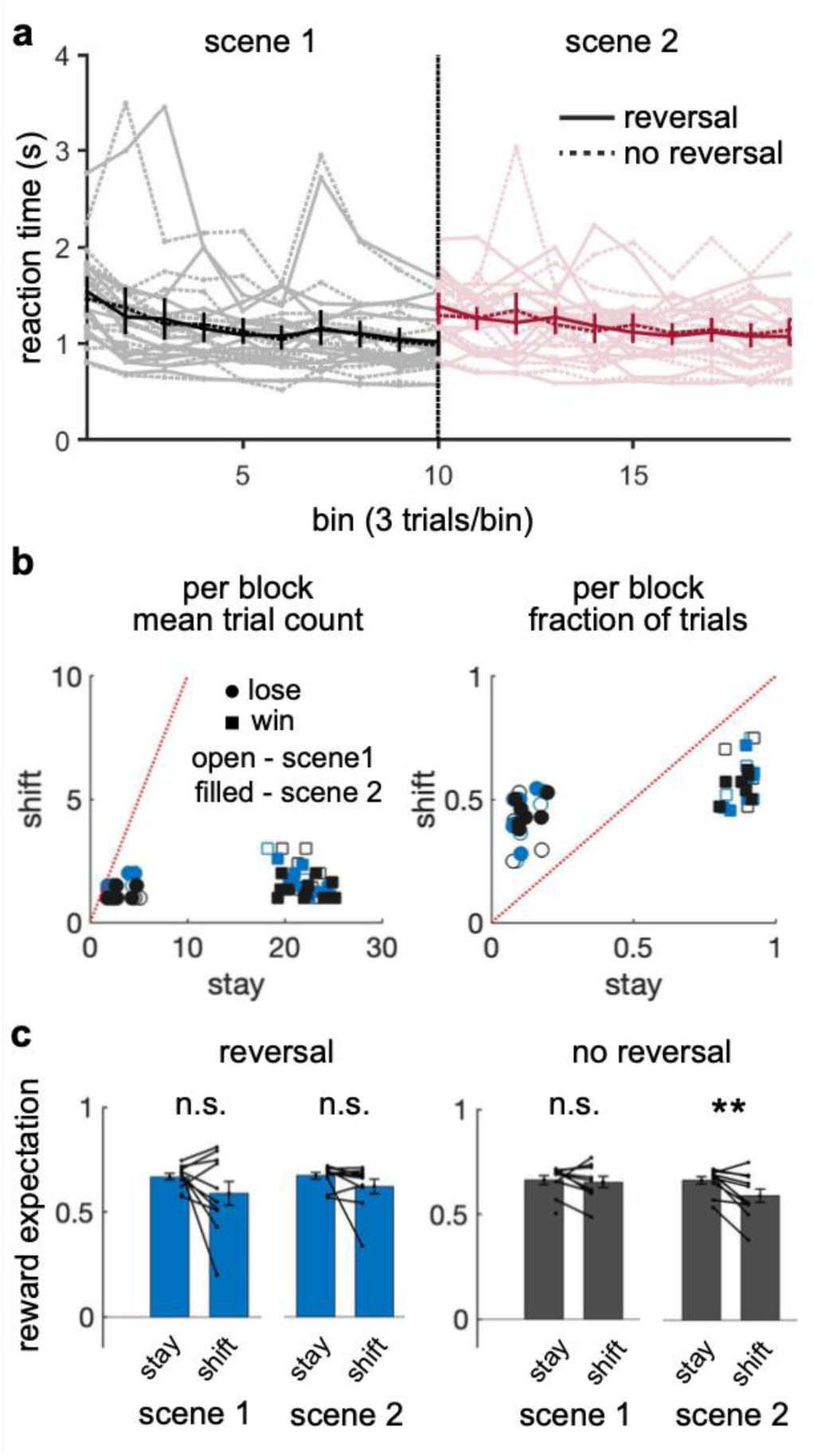
Evidence for different learning strategies between reversal and no reversal. **a**, Mean reaction times over moving bins of three trials with no overlap. Dark line and error bars show mean +/− s.e.m. across subjects. Light line shows individual subjects. **b**, Mean number of trials per block in which subjects stay or shift choices after a reward (win) or no reward (lose). Each data point is a subject. Note, the win-stay lose-shift analysis reflects the extent to which choices relate to whether a reward is received or not. **c**, Mean fraction of trials per block in which subjects stay or shift choices after a reward (win) or no reward (lose). Each data point is a subject. Stays after no reward accounts for a small fraction of stay trials while stays after reward accounts for a majority of stay trials. Shifts after no reward accounts for approximately half of shift trials and shifts after reward accounts for the other half of shift trials. **d**, Expectation for reward during trials when subjects stay (st) or shift (sh) choices after a reward is received. Mean +/− s.e.m. is shown across subjects. Significance is tested with a paired t-test across subjects (**p < .01, across subject two-tailed t-test, t(10) = 3.45, p = .0062).

**figure S3.**
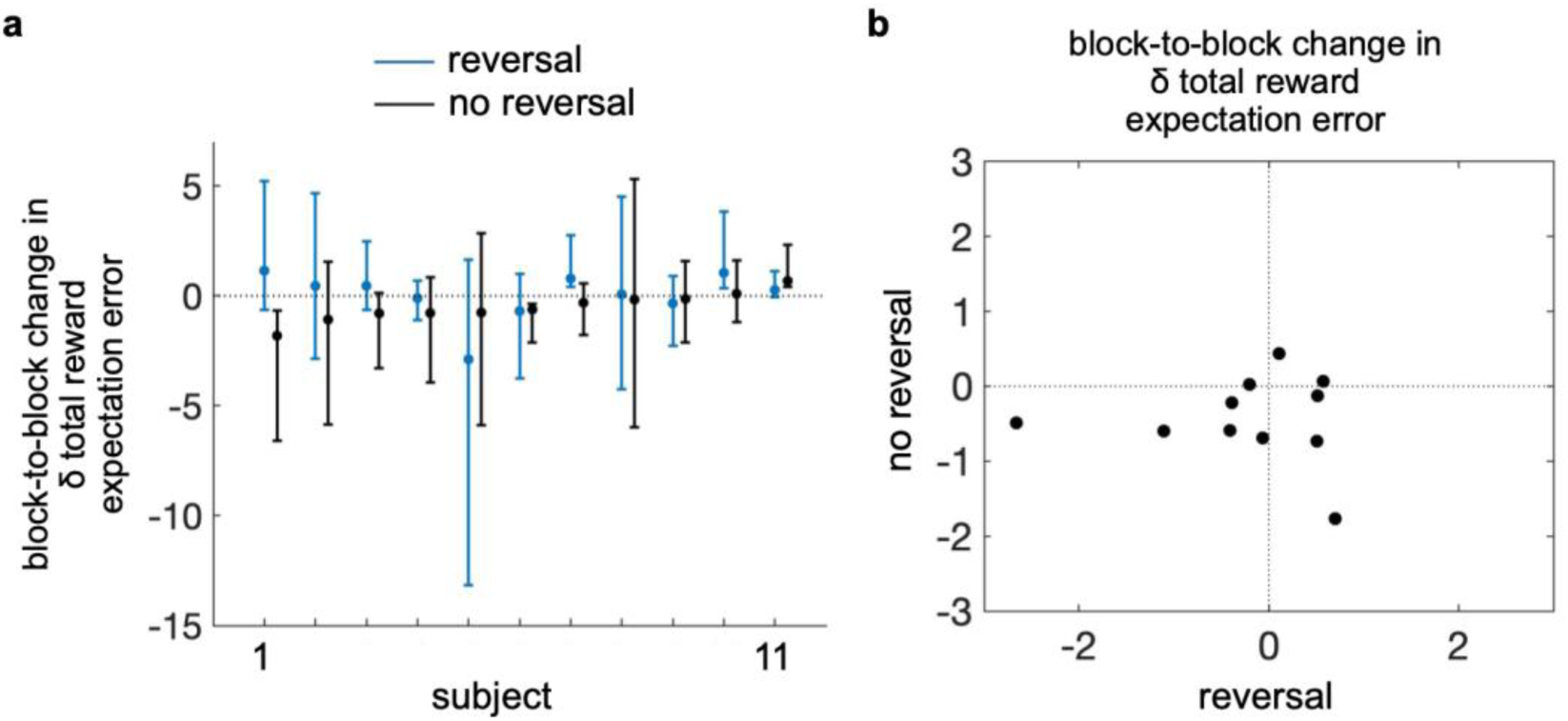
Characterization of learning efficiency. **a**, Rate of change in δ total reward expectation error between scenes as it varies over blocks for the reversal and no reversal item set for each subject, sorted by the rate for no reversal. Note, as before, δ total reward expectation can be thought of as the extent to which participants have learnt the presence of the no-reversal and reversal reward structure in the task. **b**, Comparison of rate of change in the change in total reward expectation error between scenes over blocks for reward reversal and no reversal for each subject, sorted by the rate for no reversal.

**figure S4.**
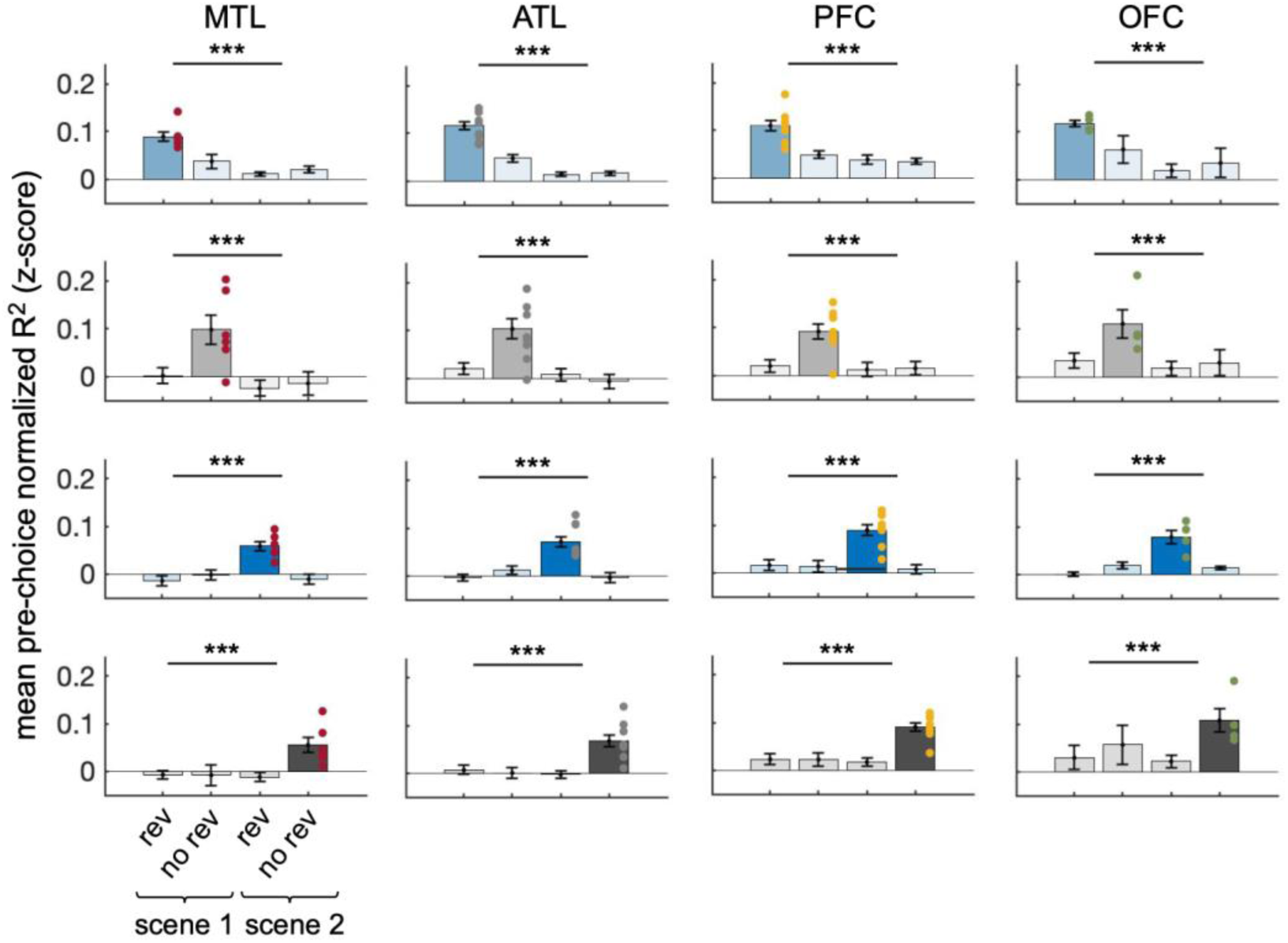
Evaluation of representations of expectation. Mean pre-choice coefficient of determination, R^2^, within and between scenes for testing if task-relevant variations are being captured, or if we are mainly getting large task-irrelevant drifts in neural power related to reward expectation. Significance in normalized R^2^ is measured by across-subject two-tailed t-tests against zero, ***p<.001. Note, the color of the bar plot refers to the trials of the item set and context for which the regression model was trained. Light colors are for scene 1 and dark colors are for scene 2; blue is for the reversal item set and black is for the no-reversal item set. The low mean pre-choice normalized R^2^ for the context conditions that are not the context condition of the neural power that was used to train the regression model indicates that the regression models are fitting to specific trial-by-trial variations for a given item set in a scene.

**figure S5.**
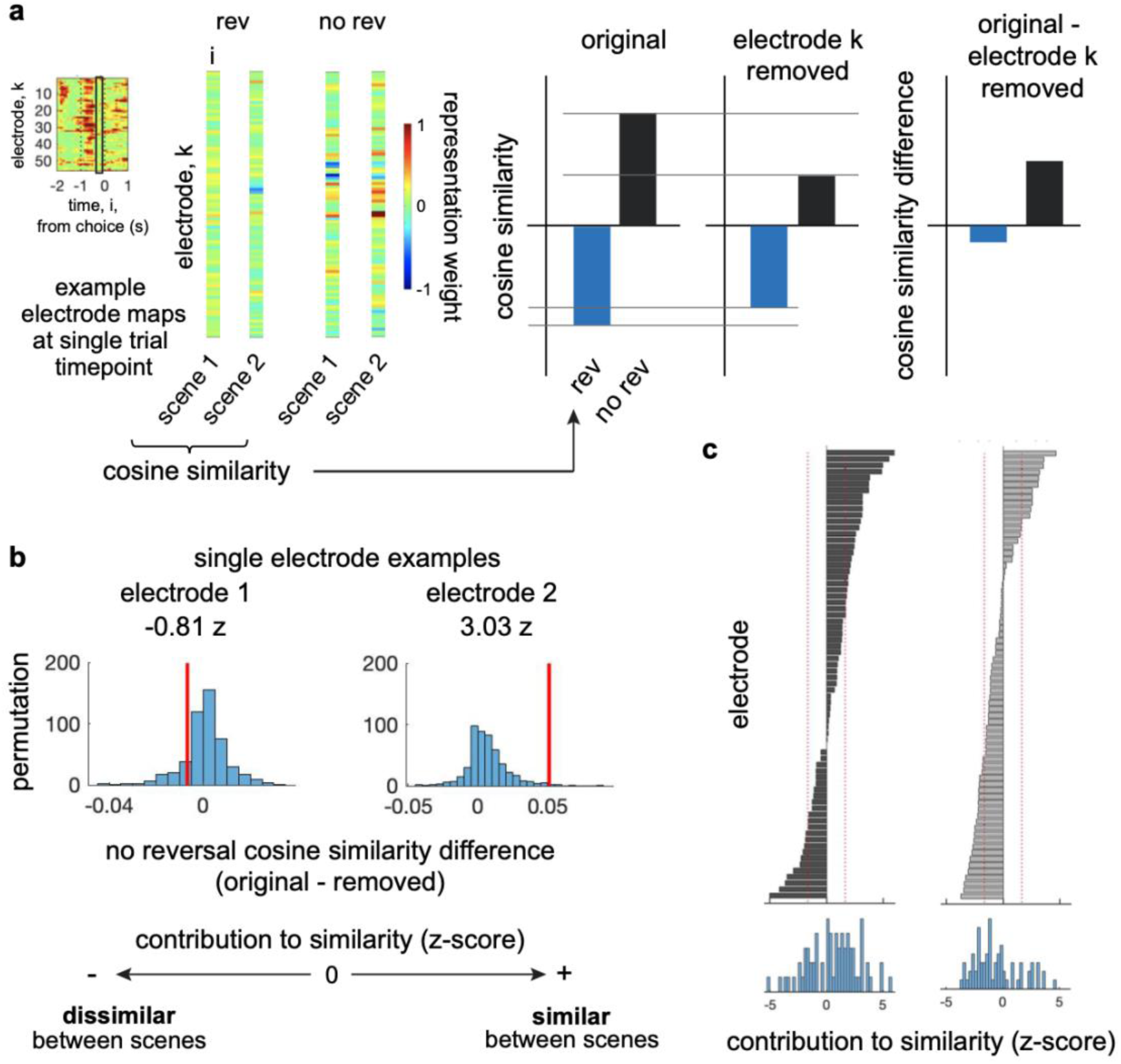
Computation of electrode-specific similarity. **a**, Schematic of cosine similarity of representation weight vectors, or electrode maps, at a single trial timepoint between scenes with all electrodes and after ablation of an electrode. **b**, Chance distribution of difference in cosine similarity with all electrodes (original) and with an electrode is ablated, and empirical difference in red. **c**, Exemplar sorted contribution of electrodes to similarity z-transformed with the mean and standard deviation of the null distribution for no reversal and reversal item sets.

**figure S6.**
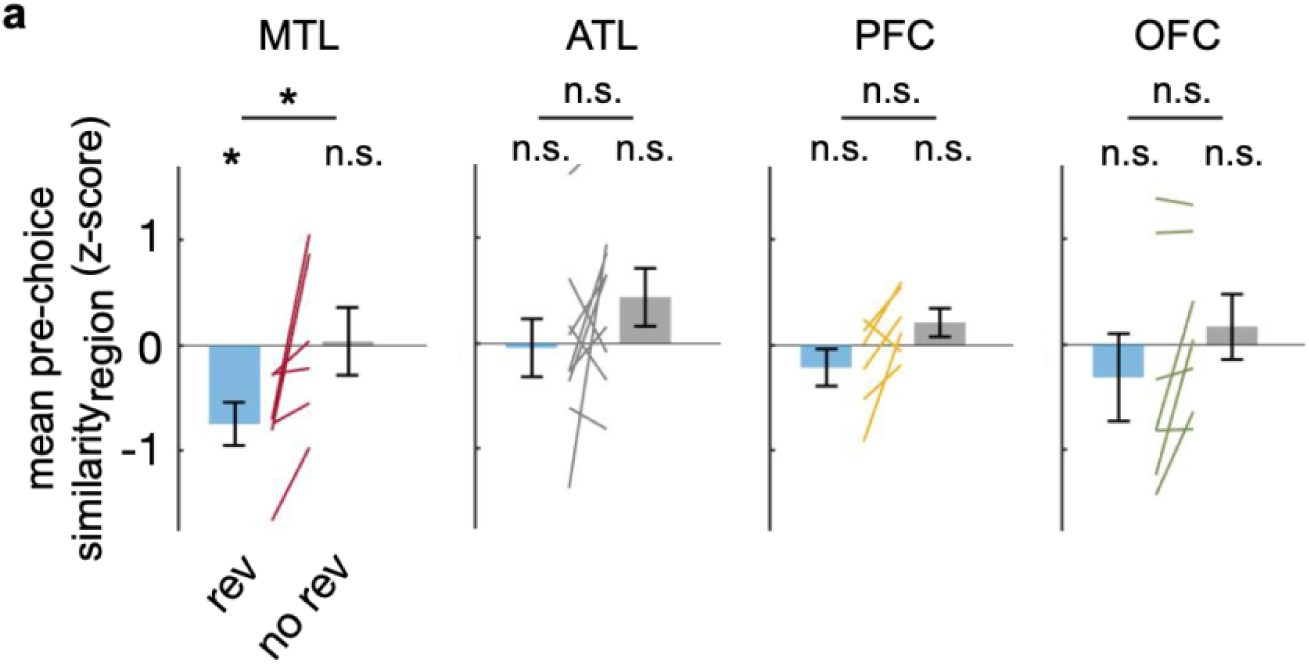
High frequency neural power is not similar between scenes. **a**, Pre-choice similarity across medial temporal lobe (MTL), anterior temporal lobe (ATL), lateral prefrontal cortex (PFC), and orbitofrontal cortex (OFC). Each line is a subject (across subject t-test; MTL: rev, t(5)=-3.63, p=.015; no rev, t(5)=0.11, p=.92; ATL: rev, t(8)=-.18, p=.86; no rev, t(8)=1.60, p=.15; PFC: rev, t(5)=-1.21, p=.28; no rev, t(5)=1.62, p=.17; OFC: rev, t(6) = –.75, p = .48; no rev, t(6)=.54, p=.61; **p<.01, n.s. p>.05).

**figure S7.**
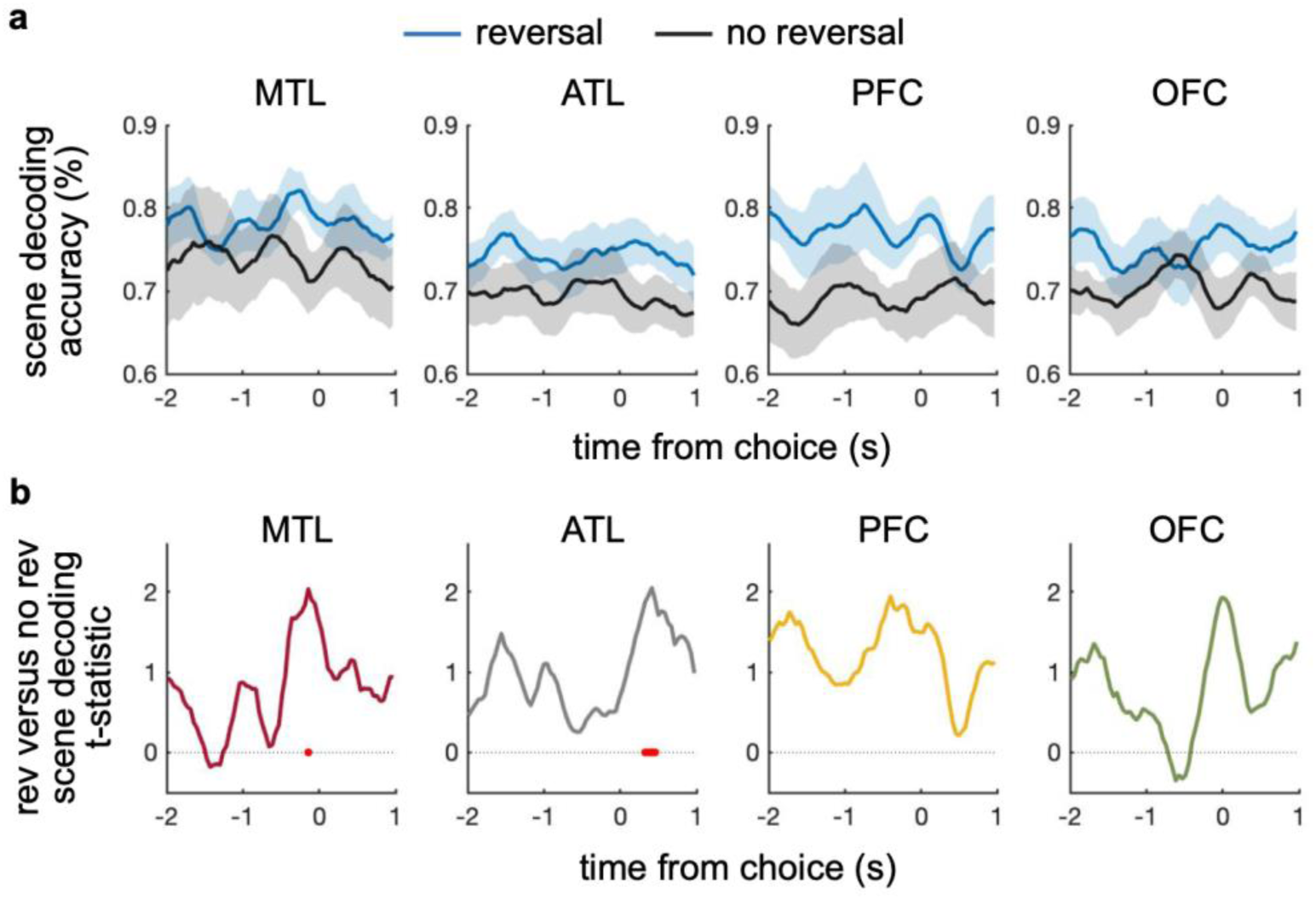
Scene decoding accuracy differs between reversal and no reversal. **a**, Accuracy of decoding scene from neural correlates of reward expectation. Shading shows the mean +/− s.e.m. across subjects. **b**, Paired t-test t-statistic of scene decoding from neural correlates of reward expectation for the reversal compared to no reversal item sets across human subjects. Red bars indicate significance p<.05.

**figure S8.**
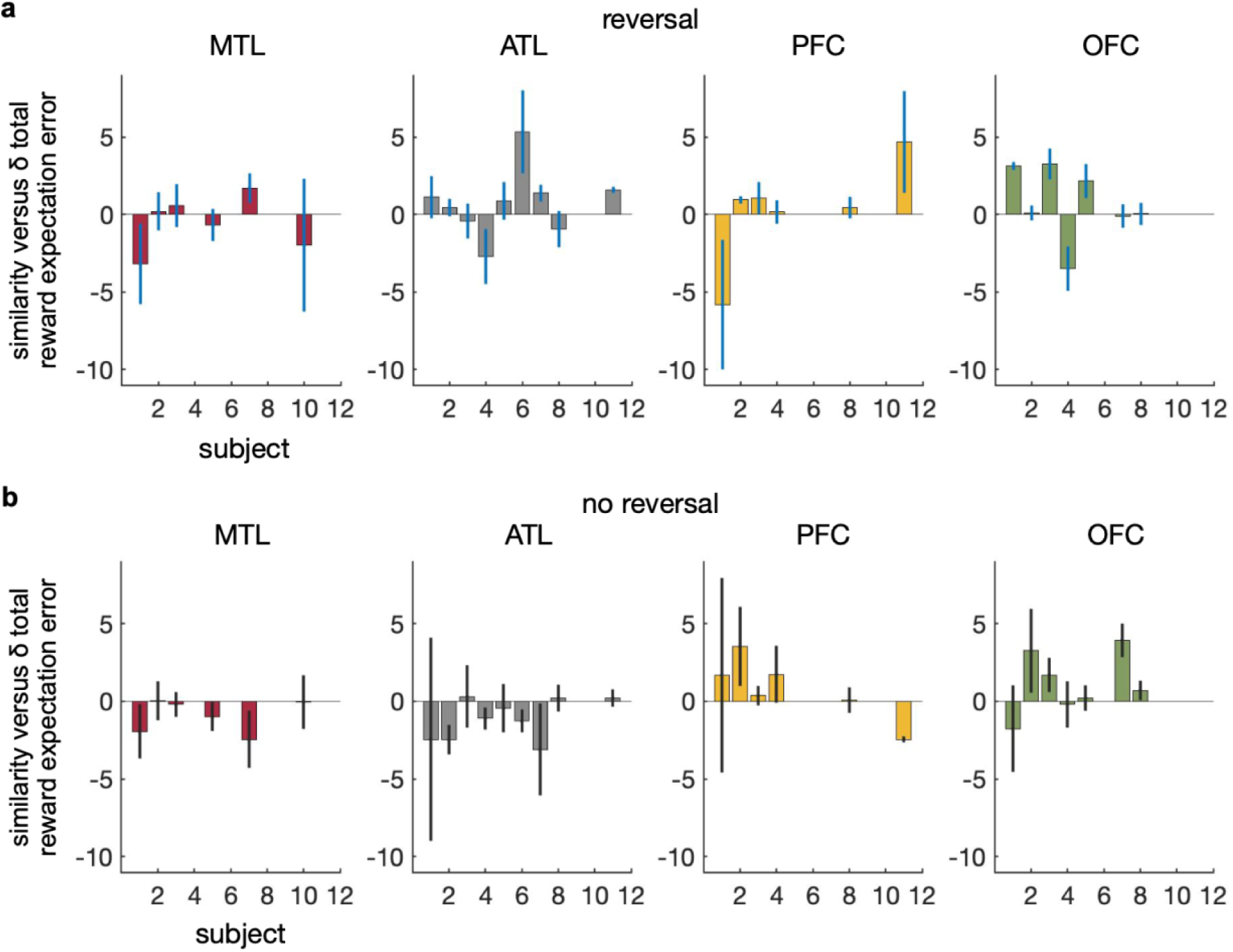
Representation similarity in ATL is related to learning when knowledge can be transferred. **a**, Regression coefficient +/− s.e.m. for each subject comparing representation similarity in the medial temporal lobe (MTL), anterior temporal lobe (ATL), prefrontal cortex (PFC), and orbitofrontal cortex (OFC) with learning, as measured by the δ total reward expectation error for the reversal item set. **b**, Regression coefficient +/− s.e.m. for each subject comparing MTL, ATL, PFC, and OFC representation similarity with learning for the no reversal item set.

**Table 1.** Patient demographics.

|  | **Age** | **Gender** | **Handed-**<br>**ness** | **Language**<br>**dominance** | **Seizure**<br>**focus** | **Resection** |
| --- | --- | --- | --- | --- | --- | --- |
| **NIH047** | 32 | male | left | bilateral | left<br>temporal<br>lobe | left temporal lobectomy,<br>amygdalohippocampectomy |
| **NIH049** | 27 | male | right | left | bilateral<br>temporal<br>lobe | none |
| **NIH050** | 20 | male | right | left | right<br>temporal<br>lobe | right temporal lobectomy,<br>amygdalohippocampectomy |
| **NIH051** | 36 | male | right | left | left<br>frontal | left cingulate topectomy |
| **NIH052** | 57 | male | right | left | right<br>temporal<br>lobe | right temporal lobectomy,<br>amygdalohippocampectomy |
| **NIH053** | 53 | female | right | left | bilateral<br>temporal<br>lobe | right temporal lobectomy,<br>amygdalohippocampectomy |
| **NIH057** | 34 | male | right | left | left<br>mesial<br>temporal<br>lobe | left anterior temporal<br>lobectomy,<br>amygdalohippocampectomy |
| **NIH058** | 22 | male | right | left | right<br>frontal<br>cortex | right frontal topectomy |
| **NIH060** | 20 | female | right | left | left<br>amygdala | selective<br>amygdalohippocampectomy |
| **NIH061** | 46 | male | right | left | left<br>medial<br>temporal<br>lobe | left selective<br>amygdalohippocampectomy |
| **NIH064** | 45 | male | right | left | right<br>temporal<br>lobe | right posterior temporal lobe<br>topectomy |

**Table 2.** Number of electrodes in brain regions of interest (MTL – medial temporal lobe; ATL – anterior temporal lobe; PFC – dorsolateral prefrontal cortex; OFC – orbitofrontal cortex; All – whole brain coverage). Total number of subjects for each brain region of interest.

|  | **MTL** | **ATL** | **PFC** | **OFC** | **All** |
| --- | --- | --- | --- | --- | --- |
| **NIH047** | 1 | 7-8 | 2 | 4 | 92-96 |
| **NIH049** | 3 | 8 | 15 | 2 | 59-64 |
| **NIH050** | 4 | 18 | 3 | 3 | 74 |
| **NIH051** |  | 1 | 28 | 15 | 104-106 |
| **NIH052** | 4-5 | 9-10 | 6 | 6 | 45-46 |
| **NIH053** |  | 15 |  |  | 56 |
| **NIH057** | 7 | 13 |  | 2 | 78-80 |
| **NIH058** |  | 1 | 29-35 | 4 | 94 |
| **NIH060** |  |  |  |  | 28 |
| **NIH061** | 1 |  |  |  | 20 |
| **NIH064** |  | 11-12 | 4 |  | 89-97 |
| **Total no. of subjects** | 6 | 9 | 7 | 7 |  |

**figure S9.**
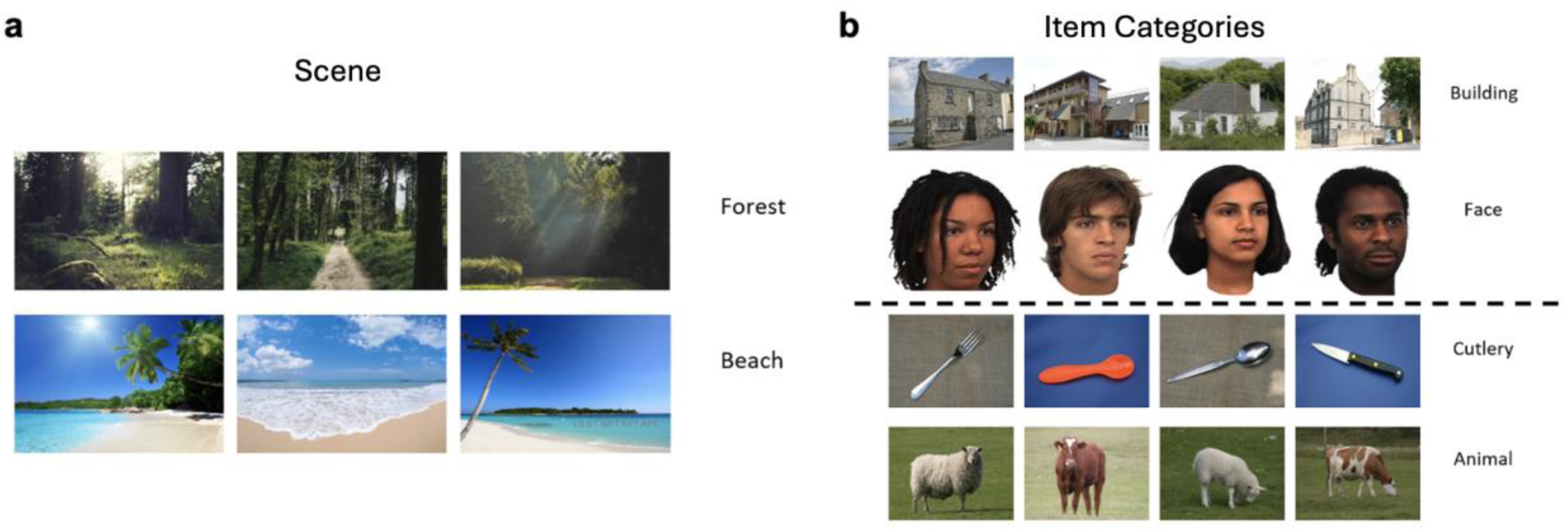
Examples of task stimuli. a, Example screens of the forest and beach context. b, Example images of items presented in each building, face, cutlery, and animal item category. The images of faces used in this study came from the Face Place dataset.^76^

**figure S10.**
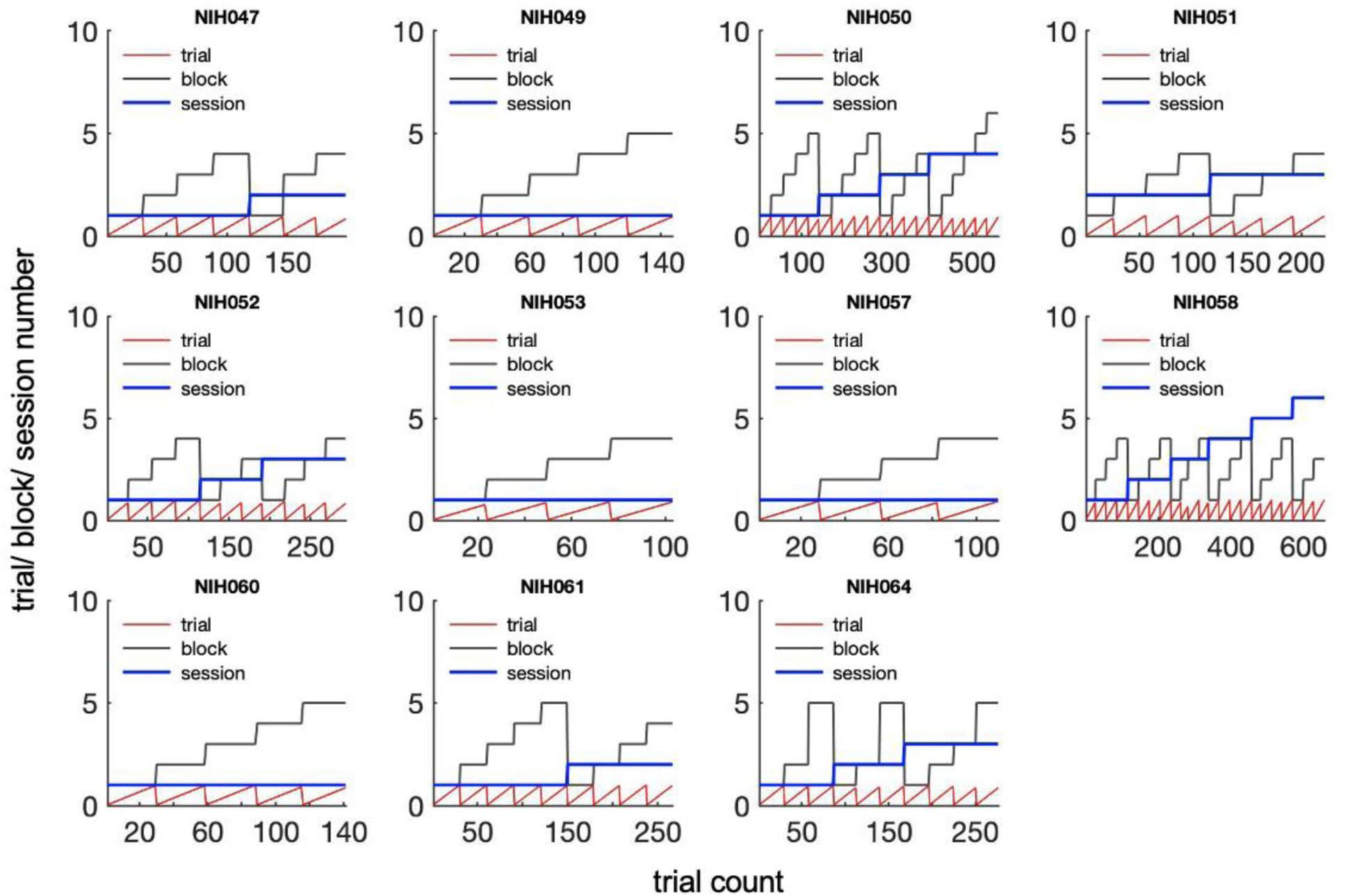
Number of trials, blocks, and sessions for each subject for one context condition.

## Supplemental information

Similar performance can be achieved by different strategies and choice patterns differ depending on strategy.^25,36,77,78^ We observed similar reaction times on average for reversal and no reversal choices. The probability that subjects chose the item with the high probability of reward does not reflect how they used current or prior reward to guide their choices. To investigate whether the expectation for reward or the reward received from the prior trial was associated with subjects’ choices (stay or shift) we measured the reward expectation related to when subjects stay or shift choices after a reward. When there is a reward reversal in scene 2, there is no difference in reward expectation between when subjects stay or shift, suggesting reward expectation is not used to guide choices. When there is no reward reversal, the expectation for reward associated with when subjects shift was significantly lower than when subjects stay, suggesting that reward expectation was associated with the choices made. This finding is expected because the reward probability was the same between scene 1 and scene 2 so it would make sense that an expectation for reward is computed and used (fig. S2).

How learning in the setting of no reversal relates to learning in the setting of a reversal is unclear. To examine the relationship, we looked at whether subjects who learned more efficiently with no reversal learned less efficiently in the setting of a reversal.

There was no linear relationship between learning efficiency in the reversal and no reversal item sets (fig. S3). While most subjects learned more efficiently when there was no reversal, the extent to which subjects learned more efficiently when there was a reversal varied across subjects.

We also examined the extent to which high frequency neural power was similar between scenes to determine whether reactivation of representations is a specific finding to variations related to reward expectation. High frequency neural power across MTL, ATL, PFC, and OFC were not significantly similar between scenes (fig. S6).

We examined the change in similarity or reactivation of representations over scene contexts in each region. There is a linear increase in similarity of representations of expectation in ATL when there is no reward reversal. We also examined the change in separation of representations over scene contexts in each region. There is a linear increase in scene decoding accuracy from representations of expectation in MTL when there is a reward reversal, suggesting the MTL separates representations when knowledge cannot be straightforwardly transferred between two scenes.

## Supplemental Methods

### Electrophysiological recordings

- Anatomic localization
- Electrophysiology artifact removal

## Anatomic localization

We localized electrodes in each subject by identifying high-intensity voxels in a post-operative CT image, which was co-registered to a pre-operative T1-weighted MRI. Electrode locations were adjusted to account for routine post-operative parenchymal shift by applying a standardized algorithm combining intraoperative photography, electrode spatial arrangement, and dural and pial surface reconstructions ^79^. Pial surfaces were reconstructed using FreeSurfer (http://surfer. nmr.mgh.harvard.edu) (Fischl, 2012) and were resampled and standardized using the AFNI SUMA package (Cox, 1996). The resulting surfaces each contained 198812 vertices per hemisphere, with vertices indexed in a standardized fashion, such that for any vertex i, the i th vertex is located in an anatomically analogous location across subjects.

We aggregated vertices from the surface reconstruction into a standard set of surface-based regions of interest (ROIs) as previously described ^79^. Briefly, we sampled 2400 equally spaced vertices per hemisphere to use as ROI centers. ROI centers were uniformly distributed across the surface at an average geodesic distance of approximately 5 mm. We assigned all vertices within a 10 mm geodesic radius of an ROI center to that ROI, which achieves a coverage of 99.9% coverage or greater of the pial surface in each subject ^79^. Because ROIs overlap, vertices may be assigned to multiple ROIs. On average, there were 669.44 ± 74.30 vertices per ROI and each vertex mapped to 8.08 ± .90 ROIs. We modeled each electrode as a cylinder with radius 1.5 mm, found the pial vertices closest to it, and then assigned each electrode to the same ROIs as its nearest pial vertices. Due to the overlap between ROIs, each electrode is assigned to multiple ROIs and each ROI may contain more than one electrode.

## Electrophysiology artifact removal

We implemented several measures to provide the most conservative sampling of non-pathological signals possible. We implemented a previously reported automated trial and electrode rejection procedure based on excessive kurtosis or variance of iEEG signals to exclude high-frequency activity associated with epileptiform activity ^74,80^ (Jang et al., 2017). We calculated and sorted the mean iEEG voltage across all trials, and divided the distribution into quartiles. We identified trial outliers by setting a threshold, Q3 + w*(Q3-Q1), where Q1 and Q3 are the mean voltage boundaries of the first and third quartiles, respectively. We empirically determined the weight w to be 2.3. We excluded all trials with mean voltage that exceeded this threshold. We visually inspected the resulting iEEG traces and found that the automated procedure reliably removed IEDs and other artifacts. In total, following exclusion of electrodes because of artifact, we retained 751 electrodes (mean ± s.e.m., 68 ± 28 electrodes per subject) for analysis.

